# B-cell SIGLEC-5 engages T-cell components of the elastin receptor complex (ERC) to suppress inflammatory T-cell cytokines

**DOI:** 10.64898/2026.06.24.734204

**Authors:** Christopher J.M. Piper, Clive Metcalfe, Maana Layeghi, Guillem Montamat-Garcia, Zara Baig, Ashley Ferrier Espósito, Lars Nitschke, Diego M Catalán, Claudia Mauri

## Abstract

SIGLECs remain poorly defined in human B-cell biology beyond SIGLEC-2/CD22 and SIGLEC-10. Here, we identify a previously unrecognized regulatory pathway involving the paired receptors SIGLEC-5 and SIGLEC-14 at the human B–T-cell interface. We show that activated B-cells differentially regulate these receptors: SIGLEC-5 is predominantly surface-expressed and induced by CD40 engagement, whereas SIGLEC-14 is primarily secreted and upregulated after both CD40 and TLR9 stimulation. We further identify EBP (elastin binding protein) and CTSA (cathepsin A) components of the elastin receptor complex (ERC), expressed by activated T-cells, as a novel ligand for both SIGLEC-5 and SIGLEC-14. Functionally, ERC-associated engagement of SIGLEC-5 on B-cells suppresses T-cell IFN-γ and IL-17 expression, establishing SIGLEC-5 as a B-cell-expressed inhibitory SIGLEC that restrains inflammatory T-cell cytokine responses. SIGLEC-14 does not alter this suppression, as SIGLEC-5□ B-cells from SIGLEC-14-sufficient and -null individuals show comparable inhibitory activity. These findings broaden SIGLEC-mediated adaptive immune regulation, with relevance to inflammatory and autoimmune disease.

## Main

The sialic acid binding immunoglobulin (Ig)-type lectin (SIGLEC) receptor family, widely expressed throughout the immune system, have gained interest as potential targetable molecules in autoimmunity and in cancer due to their ability to distinguish self from non-self. The SIGLEC family bind to sialic-acid-containing glycans commonly found on mammal glycoproteins and glycolipids, but which are typically absent on microbes. In humans, the SIGLEC family consists of 14 members, most of which contain immunoreceptor tyrosine-based inhibitory (ITIM) or ITIM-like motifs that drive negative regulation upon binding to sialo-conjugates, thus preventing undesired immune activation^1^. On the other hand, only three human SIGLECs deliver activating signals –SIGLEC-14, -15 and -16–, thanks to the recruitment of the DNAX-activating proteins of 10/12 kDa (DAP10/12) adaptors carrying an immunoreceptor tyrosine-based activating motif (ITAM)^1^. Activatory SIGLECs are typically found co-expressed with their inhibitory counterparts^1^. Given the self-recognizing nature of these receptors, tight regulation of their expression is needed to prevent the development of autoimmunity and immune escape by pathogens or cancer cells.

Human B-cells have been described to express only inhibitory SIGLECs: CD22 (SIGLEC-2), SIGLEC-5, SIGLEC-6 and SIGLEC-10^1^. CD22 and SIGLEC-10 play a pivotal role in negatively regulating B-cell antigen receptor (BCR) downstream signals^2^. Absence of CD22 predisposes autoantibody development in autoimmune-prone mouse strains, while the combined deficiency of CD22 and SIGLEC-G (the murine orthologue of SIGLEC-10) promote a lupus-like syndrome in aged mice^3^.

SIGLEC-5 has a paired activatory receptor (SIGLEC-14), which arose from gene duplication and as a result shares almost identical extracellular ligand binding domains^4^. Therefore, determination of SIGLEC-5 and SIGLEC-14 cannot be distinguished by antibodies targeting their extracellular domains. Unlike other antigen presenting cells (APCs), B-cells are known to express SIGLEC-5 and are thought not to express the polymorphic paired receptor SIGLEC-14 on the surface of the cell^5^. SIGLEC-14 is found as both membrane bound and soluble forms in other APCs, such as monocytes^6^. We aimed to characterise the expression, function and ligands of these paired receptors in B-cells.

To assess the expression of SIGLEC-5 (S5) and SIGLEC-14 (S14), we performed immunoblot analysis on B-cells isolated from SIGLEC-14^+/+^ and SIGLEC-14 null (^-/-^) individuals (these individuals lack SIGLEC-14 due to the fusion of the SIGLEC-5 and SIGLEC-14 alleles; Extended Data Fig. 1a)^5^. Both SIGLEC-5 and two distinct molecular weight forms of SIGLEC-14 were detected in B-cell lysates in SIGLEC-14^+^ individuals (Fig. 1a). To determine the cellular localisation of these receptors, we performed subcellular fractionation of healthy donor B-cells. SIGLEC-5 was detected in both membrane and cytoplasmic fractions, whereas SIGLEC-14 was predominantly confined to the cytoplasmic fraction (Fig. 1b). Consistent with this observation, pull-down of biotinylated surface proteins confirmed that only SIGLEC-5, and not SIGLEC-14, was expressed at the B-cell surface (Fig. 1c).

**Figure 1.**
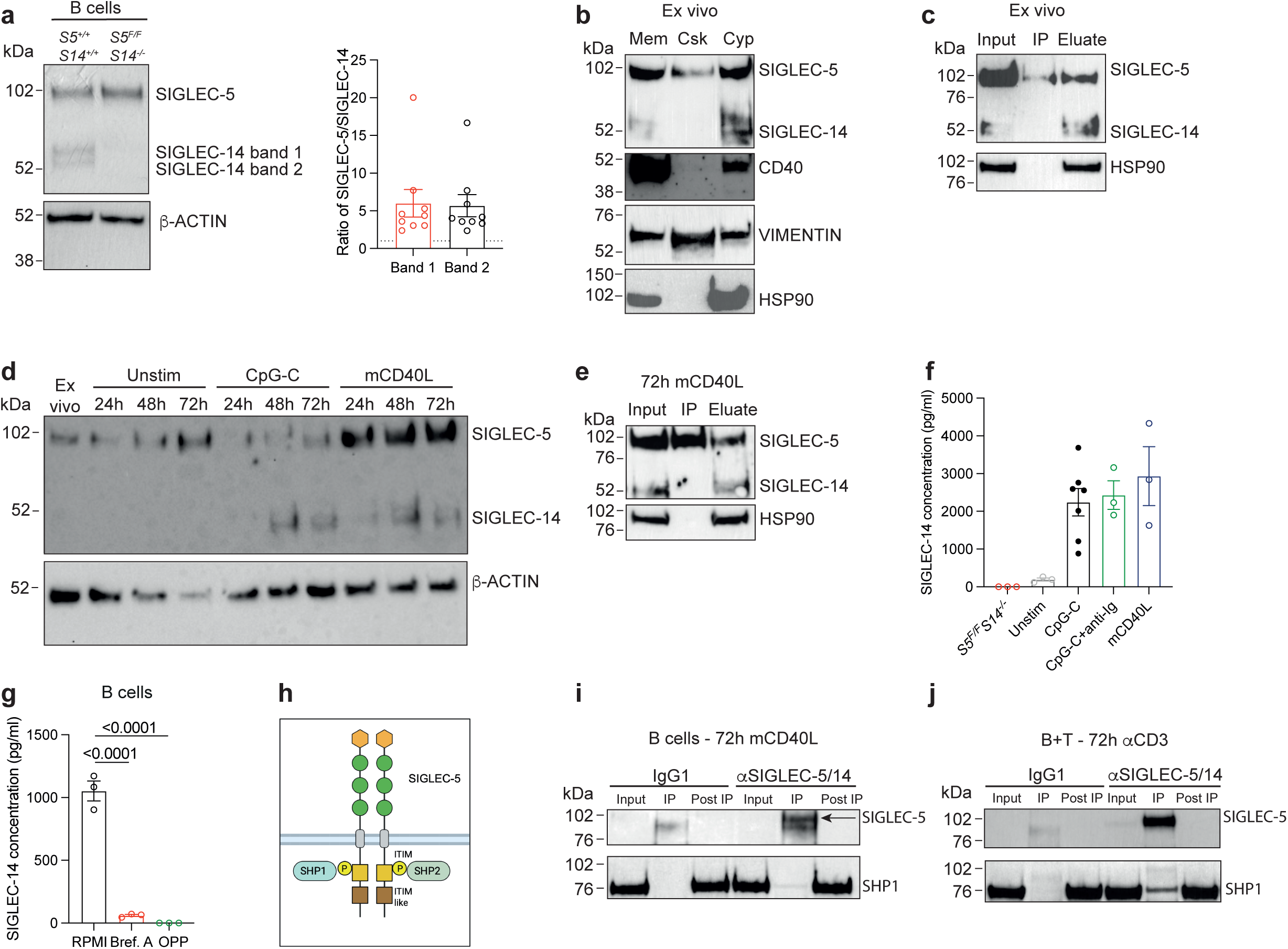
T-cells drive expression and activation of SIGLEC-5 on B-cells. **a** Immunoblots of SIGLEC-5 and SIGLEC-14 expression in B-cells from SIGLEC-14 homozygous positive (*S5^+/+^S14^+/+^*) and SIGLEC-14 null individuals (*S5^F/F^S14^-/-^*); F denotes the fusion allele between the SIGLEC-5 and SIGLEC-14 genes. Ratio of SIGLEC-5 to SIGLEC-14 band intensity in healthy individuals for both SIGLEC-14 bands (n=9). **b,** SIGLEC-14^+/+^ B-cells were split into sub-cellular fractions and assessed by immunoblots for SIGLEC-5, SIGLEC-14, CD40, VIMENTIN and HSP90. **c,** Immunoblots of SIGLEC-5 and SIGLEC-14 after pull-down of biotinylated ex vivo B-cell surface proteins. **d,** B-cells were isolated from healthy controls and either left unstimulated or stimulated with 1μM CpG-C or 1μg/ml mega-CD40L for the indicated time points. The lysates were assessed by immunoblot for SIGLEC-5 and SIGLEC-14. **e,** Immunoblots of SIGLEC-5 and SIGLEC-14 after pull-down of biotinylated surface proteins from 72h mCD40L stimulated B-cells. **f,** Supernatants were taken from CD40L stimulated SIGLEC-14 null (*S5^F/F^S14^-/-^*) B-cells and CpG-C, CpG-C+anti-Ig or CD40L stimulated SIGLEC-14^+^ B-cells and were assessed for SIGLEC-14 expression by ELISA (n=3 and n=6). **g,** B-cells were activated for 24h with 1μg/ml megaCD40L and then incubated for a further 18h in the presence of either Brefeldin A (5μg/ml) or O-propargyl-puromycin (OPP; 50μM). Supernatants were assessed for the levels of SIGLEC-14 by ELISA (n=3). **h**, Schematic illustrating the SIGLEC-5 signalling pathway. **i,** Immunoblots of SIGLEC-5 and SHP1 in 72h mCD40L stimulated B-cells, after immunoprecipitation with a IgG1 control antibody or a SIGLEC-5 antibody (1A5). **j,** Immunoblots of SIGLEC-5 and SHP1 in 72h αCD3 stimulated CD4^+^ T-cell and B-cell co-cultures, after immunoprecipitation with a IgG1 control antibody or a SIGLEC5 antibody (1A5). For immunoblots, β-ACTIN was used as a loading control. For subcellular fractions CD40, VIMENTIN and HSP90 were used as fractionation controls (membrane, cytoskeletal and cytoplasmic respectively). All data are representative of at least two independent experiments. All values represent the mean ± SEM; one-way ANOVA for **g**.

SIGLEC-14 has been described in other APCs in both membrane-bound and soluble forms^6^. As membrane-bound SIGLEC-14 requires DAP12 for signalling^4^, we assessed DAP12 expression as a proxy for a functional indicator of surface-associated SIGLEC-14. Unlike monocytes, which co-express SIGLEC-5 (Extended Data Fig. 1b-c, e), SIGLEC-14 (Extended Data Fig. 1d-e) and DAP12 (Extended Data Fig. 1 f-g), B-cells lacked detectable DAP12 (Extended Data Fig. 1f).

Given that SIGLEC oligomerisation may influence their inhibitory function, next we investigated whether SIGLEC-5 adopts a homodimeric configuration in B-cells, as has been reported in monocytes^7^. Non-denaturing immunoblot analysis of lysates from SIGLEC-14-null B-cells revealed that, similarly to myeloid cells, SIGLEC-5 exists as a homodimer (Extended Data Fig. 2a). Having established that only SIGLEC-5, of the SIGLEC-5/SIGLEC-14 receptor pair, is expressed on the surface of B-cells (Fig. 1b-c, e), next, we assessed its distribution across B-cell subsets. Flow cytometric analysis demonstrated a maturation-associated increase in SIGLEC-5 expression, with the highest levels observed in unswitched memory B-cells (Extended Data Fig. 2b,c).

To investigate whether distinct activation pathways differentially regulate SIGLEC-5 and SIGLEC-14 expression, we analysed the kinetics of both receptors in B-cells in response to T-cell–dependent (CD40 ligand (CD40L)) and independent (TLR9, CpG-C) stimulation. Compared to the *ex vivo* condition, we observed increased expression of SIGLEC-5 after activation with CD40L, whilst stimulation with CpG-C downregulated total protein levels of SIGLEC-5 (Fig. 1d). Pull-down of biotinylated cell surface proteins from CD40L activated B-cells confirmed that, like in *ex vivo* B-cells, SIGLEC-5 expression, and not SIGLEC-14, is expressed at the cell surface (Fig. 1e).

SIGLEC-14 was expressed at lower levels than SIGLEC-5 and was virtually absent in unstimulated B-cells, but increased after activation through TLR9, TLR9+BCR and CD40 pathways (Fig. 1d,f). Unlike SIGLEC-5, which was detected at the plasma membrane, SIGLEC-14 was confined to the cytoplasmic fraction and was not detected at the B-cell surface (Fig. 1c,e). To test whether B-cell activation promotes the release of SIGLEC-14 and/or SIGLEC-5 as a soluble protein after activation, we purified B-cells from SIGLEC-5/14-sufficient and SIGLEC-14-null individuals. Following CD40L stimulation, B-cells from SIGLEC-5/14-sufficient individuals released a detectable soluble SIGLEC-5/14 protein, whereas CD40L-activated B-cells from SIGLEC-14-null individuals did not released any detectable SIGLEC-5 or 14 protein. This demonstrates that the soluble protein detected after activation was SIGLEC-14 and not SIGLEC-5. SIGLEC-14 was also secreted upon CpG-C stimulation (Fig. 1f).

As soluble SIGLEC-14 has been described in monocytes^6^, we compared the kinetics of SIGLEC-14 secretion between monocytes and B-cells. Whereas, B-cells required activation to secrete SIGLEC-14 (Fig. 1d,f), monocytes secreted SIGLEC-14 even in the absence of activation (Extended Data Fig. 3a). Similarly to monocytes^6^, B-cells also secreted SIGLEC-14 as a homodimer (Extended Data Fig. 3b).

To determine whether soluble SIGLEC-14 resulted from active secretion rather than proteolytic shedding, we blocked protein transport with Brefeldin A or inhibited translation using O-propargyl-puromycin. Both treatments markedly reduced SIGLEC-14 levels in the supernatant of activated B-cells, suggesting SIGLEC-14 is actively secreted (Fig. 1g). Thus, we propose that activated human B-cells display a clear division in SIGLEC-5 and SIGLEC-14 biology: SIGLEC-5 functions as a surface-expressed receptor, whose expression is upregulated by CD40 signalling, whereas SIGLEC-14 is released as a soluble molecule in response to CD40 and TLR9 engagement.

SIGLEC-5 and SIGLEC-14 bind a broad range of sialylated and non-sialylated ligands, including DAMPs^4, 8, 9, 10^, and can engage ligands either in *cis* (on the same cell membrane) or in *trans* (e.g. on adjacent T-cells or soluble molecules)^1^. Engagement of SIGLEC-5 by sialylated ligands presented in *trans* induces phosphorylation of its ITIM motifs and recruitment of SHP1/SHP2 phosphatases^11^ (Fig. 1h), consistent with a model in which neighbouring T-cells regulate B–T-cell cross-talk through interaction with SIGLEC-5. This possibility is further supported by the observation that SIGLEC-5 expression is upregulated during B–T-cell interactions (Fig. 1d). We firstly asked whether contact with activated T-cells is sufficient to promote SIGLEC-5 signalling in B-cells.

To determine whether SIGLEC-5 is functionally engaged during B–T-cell interactions, we assessed SHP1 association following receptor immunoprecipitation. The absence of surface SIGLEC-14, together with its release as a soluble homodimer, confined this contact-dependent signalling analysis to membrane-expressed SIGLEC-5. CD40L alone induced minimal SHP1 recruitment to SIGLEC-5 (Fig. 1i), indicating that SIGLEC-5 is not activated in *cis* under these conditions. By contrast, anti-CD3–stimulated CD4□ T-cells induced SHP1 recruitment to B-cell-derived SIGLEC-5 (Fig. 1j). As anti-CD3/CD28–stimulated CD4□ T-cells did not express SIGLEC-5 in our system, despite previous reports¹², this excluded T-cell-derived SIGLEC-5 as the immunoprecipitated receptor (Extended Data Fig. 3c), thus demonstrating that activated T-cells functionally engage B-cell SIGLEC-5 *in trans*.

The endogenous ligands of SIGLEC-5 and SIGLEC-14 in the context of B-cell biology remain unknown. As interaction with activated T-cells upregulated SIGLEC-5 on B-cells, next we hypothesised that activated T-cells express ligands capable of engaging the SIGLEC-5/SIGLEC-14 receptor pair. To identify putative T-cell ligands for B-cell membrane-bound SIGLEC-5 and secreted SIGLEC-14, anti-CD3–stimulated CD4□ T-cells were incubated with recombinant human IgG1-Fc, SIGLEC-5-Fc or SIGLEC-14-Fc, followed by pulldown using protein G Dynabeads and LC-MS/MS analysis (Extended Data Fig. 3d). Silver staining and immunoblotting confirmed successful enrichment of T-cell–derived binding partners (Fig. 2a). Notably, greater amounts of SIGLEC-14-Fc were recovered compared to SIGLEC-5-Fc, suggesting higher affinity or increased ligand availability (Fig. 2a).

**Figure 2.**
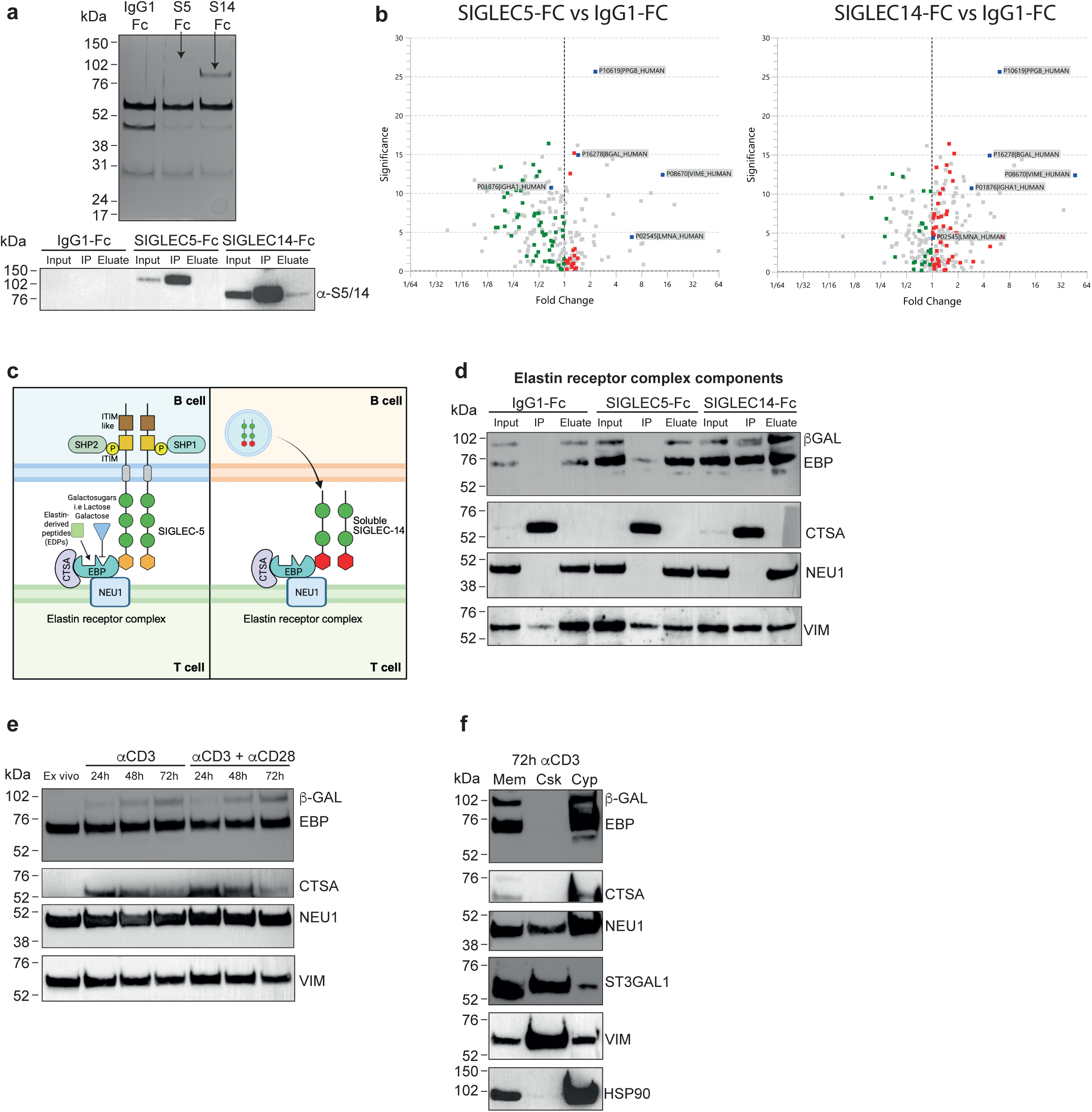
Identification of the elastin binding protein (EBP) as a ligand of SIGLEC-5 and SIGLEC-14. **a-b**, CD4^+^ T-cells were tagged with recombinant IgG1-Fc, SIGLEC5-Fc (S5-Fc) or SIGLEC14-Fc (S14-Fc) and associated ligands were immunoprecipitated by protein G bound beads. The resulting pull-downs were analysed by LC-MS/MS, silver staining or immunoprecipitation. **a,** Silver-stained gel of IgG1-Fc, SIGLEC5-Fc and SIGLEC14-Fc tagged immunoprecipitated CD4^+^ T-cell proteins (top blot). Immunoblot of SIGLEC-5 and SIGLEC-14 on the SIGLEC5-Fc and SIGLEC14-Fc immunoprecipitates as above (bottom blot). **b,** Volcano plots of protein pulldown enrichment in IgG1-Fc versus SIGLEC5-Fc (left plot) or IgG1-Fc versus SIGLEC14-Fc (right plot) comparisons. **c,** Schematic illustrating a model for the interaction between SIGLEC-5 or SIGLEC-14 and elastin receptor complex (ERC) components: elastin-binding protein (EBP), cathepsin-A (CTSA), and neuroaminidase-1 (NEU1). **d,** Immunoblots of EBP/β-Galatosidase (β-Gal), CTSA, NEU1 and Vimentin (VIM) in IgG1-Fc, SIGLEC5-Fc and SIGLEC14-Fc tagged CD4^+^ T-cell immunoprecipitates. **e,** Immunoblots of EBP/β-GAL, CTSA, NEU1 and VIM in 72h αCD3±aCD28 stimulated CD4^+^ T-cells. **f,** CD4^+^ T-cells were stimulated for 72h with αCD3, split into sub-cellular fractionations and assessed by immunoblots for EBP/β-GAL, CTSA, NEU1 and the fraction controls ST3GAL1 (membrane), VIM (cytoskeletal) and HSP90 (cytoplasmic). All data are representative of at least two independent experiments. For the LC-MS/MS experiments, each comparison was carried out in biological triplicates, with technical triplicates for each condition. The fold change for the protein pulldown comparisons is based on label free quantification (LFQ) intensities. For immunoblots, VIMENTIN was used as a loading control.

Label-free quantification identified 49 proteins enriched in both SIGLEC-5 and SIGLEC-14 pulldowns relative to IgG1-Fc controls (Fig. 2b; Supplementary Table 1). Among these was the previously described SIGLEC-14 ligand vimentin^9^. Of particular interest, cathepsin A (CTSA) and β-galactosidase (β-GAL) were significantly enriched, both components of the elastin receptor complex (ERC) (Fig. 2b,c)^12, 13^.

β-GAL has two splice variants: a lysosomal catalytically active form and a truncated catalytically inactive membrane-associated form (elastin binding protein, EBP). The latter forms part of the elastin receptor complex (ERC), which binds elastin-derived peptides (EDPs) generated during elastin degradation (Fig. 2c)^13^. Both splice variants associate with CTSA and neuraminidase 1 (NEU1), forming a multi-protein complex. Importantly, NEU1 functions as a sialidase that removes terminal sialic acids from glycoproteins and glycolipids, potentially regulating the availability of SIGLEC ligands at the cell surface (Fig. 2c)^13^. Due to the intrinsic link between the ERC and sialic acid bioavailability, we chose to focus on the role of ERC-SIGLEC interactions on the functional outcomes of B and T-cells.

Independent immunoprecipitations confirmed that both SIGLEC-5 and SIGLEC-14 predominantly bound the EBP splice variant of β-GAL (Fig. 2d). Although CTSA was enriched in SIGLEC pulldowns, background binding was observed in IgG1-Fc controls. Neither SIGLEC directly associated with NEU1 (Fig. 2d). Consistent with these findings, EBP was the dominant β-GAL splice variant expressed in CD4□ T-cells (Fig. 2e). Importantly, activation altered ERC component expression. EBP and NEU1 levels did not change upon T-cell activation, whereas CTSA expression was promptly induced following activation but declined by 72 hours (Fig. 2e), suggesting that CTSA is the rate limiting component of the complex. Subcellular fractionation of anti-CD3–activated CD4^+^ T-cells demonstrated that EBP/β-GAL, CTSA and NEU1 were present in both membrane and cytoplasmic fractions, indicating the presence of both lysosomal and membrane-associated complexes (Fig. 2f).

To determine whether the expression of the gene components constituting the ERC complex (*EBP*, *CTSA* and *NEU1*) were associated with specific T-cell subsets following activation, we analysed a published single-cell RNA sequencing dataset of anti-CD3/CD28–activated human T-cells^14^. An ERC module score, created by averaging the expression of the ERC subunit genes (*CTSA*, *GLB1*, *NEU1*), revealed highest expression of the complex within activated CD4^+^T-cell clusters. Activated CD4^+^ T-cell clusters also displayed elevated activation markers (*CD40LG, IFNG, TNF, TNFRSF4*) and cell-cycle genes (Fig. 3a–g). Collectively, these data suggested the ERC (particularly the EBP splice variant) as a ligand for SIGLEC-5 and SIGLEC-14 and indicate that activated T-cells are enriched for ERC expression, supporting their role as a source of SIGLEC ligands during B–T-cell interactions.

**Figure 3.**
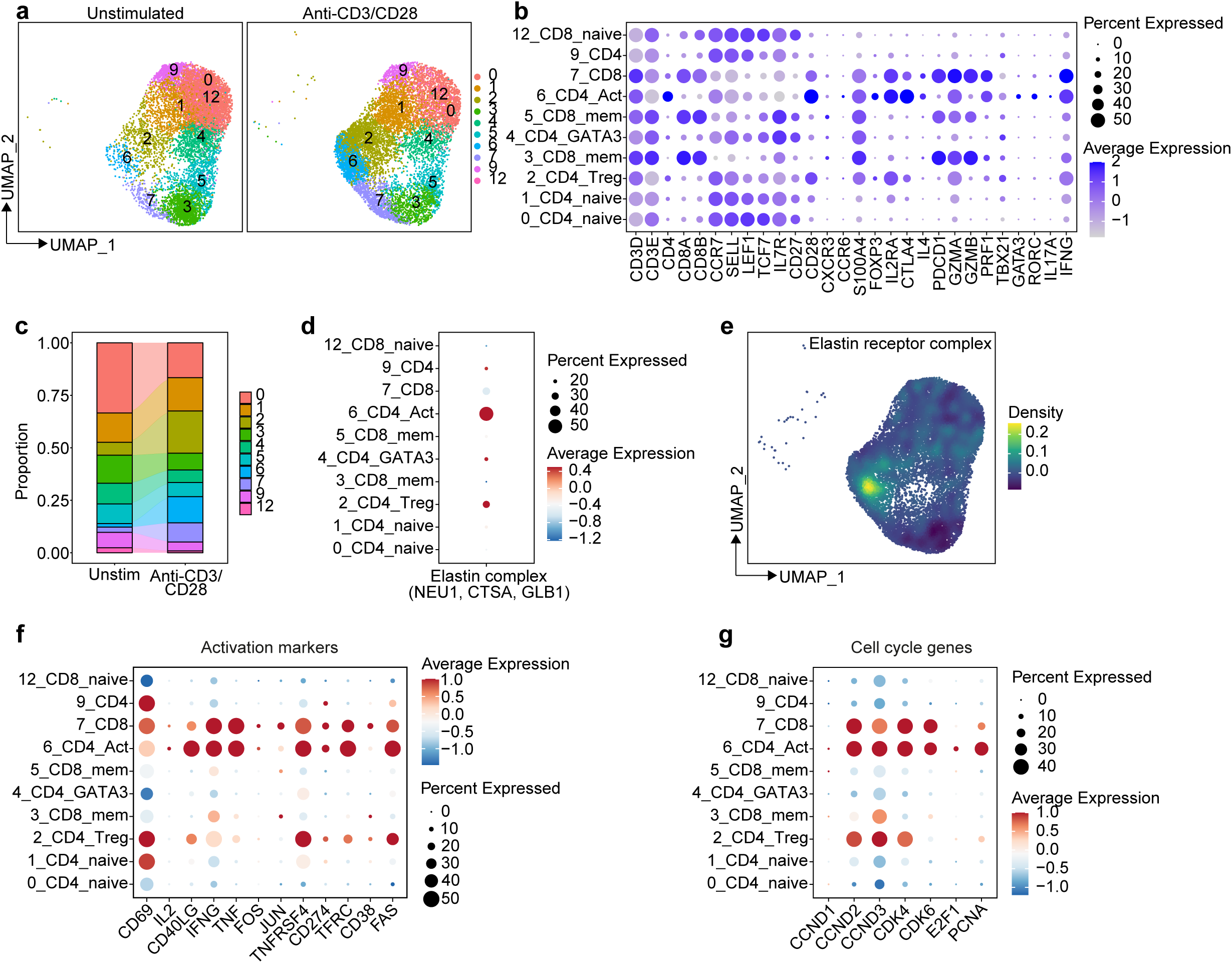
The elastin receptor complex (ERC) is highly expressed by activated CD4^+^ T-cells. **a,** UMAP showing T-cell subsets in unstimulated and αCD3+αCD28 stimulated CD3^+^ T-cells. **b,** Dot plot showing the proportion of cells expressing key T-cell lineage genes for each T-cell cluster. **c,** Stacked bar chart showing the proportion of each T-cell cluster in unstimulated and αCD3/CD28-stimulated CD3^+^ T-cells. **d,** Elastin receptor complex (ERC) score comprising *NEU1, CTSA* and *GLB1* and their average expression levels (color intensity) across T-cell clusters in αCD3/CD28-stimulated CD3^+^ T-cells. **e,** Density UMAP showing the projection of the ERC score onto the T-cell clusters of αCD3/CD28–stimulated CD3^+^ T-cells. **f,** Dot plot showing the expression of the indicated activation marker genes across the T-cell clusters. **g,** Dot plot showing the expression of the indicated cell cycle and proliferation associated genes across the T-cell clusters.

Having identified T-cell-derived SIGLEC-5 ligands, we next asked whether the outcome of activated T-cell-SIGLEC-5^+^B-cell interactions result in the inhibition of T-cell IFN-γ and IL-17 responses and whether this is EBP and CTSA dependent. We employed FACS-sorted SIGLEC-5□ and SIGLEC-5□ B-cells and co-cultured them with CD25□CD4□ T-cells to assess suppression of inflammatory cytokine responses (gating strategy; Extended Data Fig. 4a). SIGLEC-5□ B-cells, but not SIGLEC-5□ B-cells, significantly suppressed T-cell IFN-γ and IL-17 expression (Fig. 4a–c), identifying SIGLEC-5 as a functional marker of B-cell-mediated control of effector T-cell cytokines. This suppressive activity was significantly diminished after blockade of either SIGLEC-5 or the IL-10/IL-10R pathway, demonstrating that SIGLEC-5□ B-cells restrain inflammatory T-cell responses through a SIGLEC-5-dependent, IL-10-linked mechanism (Fig. 4a–c).

**Figure 4.**
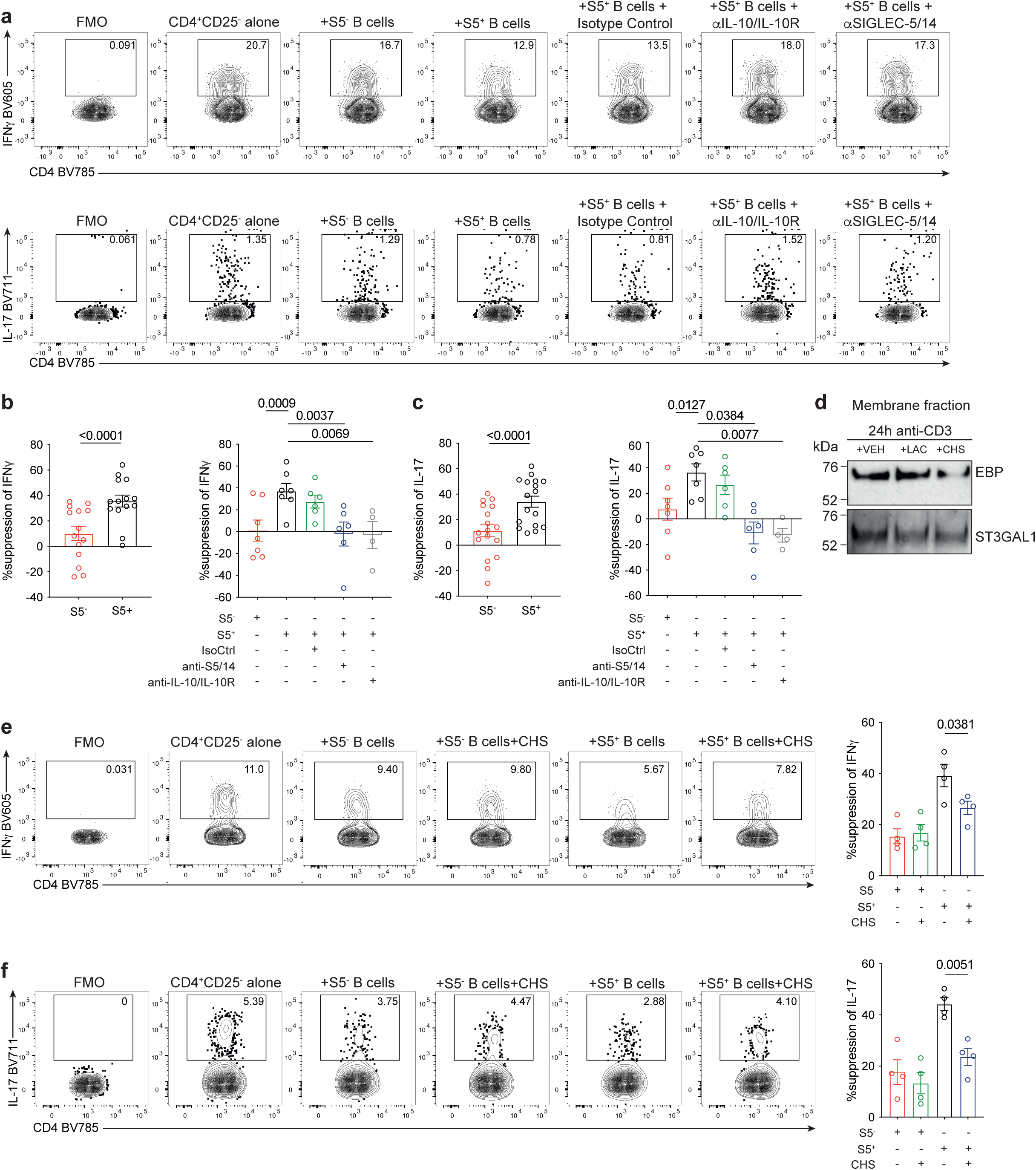
SIGLEC-5^+^ B-cell mediated suppression of T-cell IFNγ and IL-17 production is dependent on interaction with T-cell EBP. **a-c,** B-cells from *S5^+/+^S14^+/+^*individuals were stimulated for 72h with 1μM CpG-C and SIGLEC-5^+^ and SIGLEC-5^-^ B-cells were FACS sorted and co-cultured with CD4^+^CD25^-^T-cells with 0.5μg/ml plate bound anti-CD3 and CpG-C, either alone or with blocking antibodies against SIGLEC-5/14 (5μg/ml), IL-10/IL-10R (5μg/ml) or the respective IgG1 isotype control (5μg/ml). Representative flow cytometry plots and bar charts showing the percentage suppression of T-cell **a,b** IFNγ or **a,c,** IL-17 expression after the addition of SIGLEC-5^+^ and SIGLEC-5^-^ subsets (n=14 or n=17 for IFNγ and IL-17 respectively; for comparisons with blocking antibodies n=4 or more). **d,** Immunoblots for EBP and ST3GAL1 expression in the membrane fraction of 24h anti-CD3–stimulated CD4^+^ T-cells incubated with 1mM lactose (LAC), 100μM chondroitin sulphate (CHS), or a vehicle (VEH) control. ST3-β-galactoside-α-2,3-sialyltransferase-1 (ST3GAL1) was included as membrane-expressed control. **e-f,** SIGLEC-5^+^ and SIGLEC-5^-^ B-cells were FACS sorted from *S5^+/+^S14^+/+^* individuals and co-cultured with anti-CD3+CpG-C stimulated CD4^+^CD25^-^ T-cells with/without chondroitin sulphate (CHS). Representative flow cytometry plots and bar charts showing the percentage suppression of T-cell **e,** IFNγ (n=4) and **f**, IL-17 expression (n=4). Percentage suppression was calculated compared to anti-CD3+CpG-C stimulated CD4^+^CD25^-^ T-cells alone. All data are representative of at least two independent experiments. All values represent the mean ± SEM; Student’s t test for **b,c;** mixed effects analysis (Dunnett’s multiple comparison post-hoc test) for **b,c** and one-way ANOVA (Tukey’s multiple comparison post-hoc test) for **e,f**. FMO: fluorescence minus one.

To define the contribution of soluble SIGLEC-14 to B-cell-mediated suppression of T-cell responses, we isolated SIGLEC-5□ B-cells from SIGLEC-14-null individuals, who lack secreted SIGLEC-14 (Fig. 1f). SIGLEC-5□ B-cells from SIGLEC-14-null donors suppressed T-cell IFN-γ and IL-17 expression to a similar extent as SIGLEC-5□ B-cells from SIGLEC-14-sufficient donors (Extended Data Fig. 4b,c). Consistent with this result, addition of recombinant human SIGLEC-14 to anti-CD3-activated CD4□ T-cells did not alter IFN-γ or IL-17 expression (Extended Data Fig. 5a,b). Together, these data show that soluble SIGLEC-14 is dispensable for this effect and identify SIGLEC-5 as the dominant SIGLEC component through which B-cells suppress T-cell IFN-γ and IL-17 expression.

Next we assessed if SIGLEC-5 mediated suppression of T-cell responses was due to interactions with the T-cell EBP-complex, T-cells were cultured with two EBP inhibitors; lactose and chondroitin sulphate (CHS). Lactose, a galacto-sugar ligand of EBP, interferes with EBP-dependent recognition of elastin-derived ligands^15^, whilst CHS causes shedding of EBP from the cell surface leading to ERC disruption^16, 17^. Incubation of anti-CD3-stimulated T-cells with CHS, but not lactose, decreased EBP levels (Fig. 4d). In the absence of T-cell EBP, after incubation with CHS, SIGLEC-5^+^ B-cells exhibited reduced capacity to suppress CD4^+^ T-cell IFNγ and IL-17 expression (Fig. 4e,f). No differences in B-cell-mediated suppression of T-cell cytokine expression were observed after inhibition of EBP with lactose (Extended Data Fig 5c,d). Together, these data identify B-cell-expressed SIGLEC-5 as a co-inhibitory molecule involved in B-cell-mediated suppression of inflammatory T-cell IFN-γ and IL-17 responses.

SIGLEC-5 is a member of the CD33-related SIGLEC family, for which no orthologue exists in rodents. Unlike other SIGLECs, its extracellular V-type domain displays a broad binding preference for α2,3-, α2,6- and α2,8-linked sialic acids^18^. In myeloid cells, SIGLEC-5 has been described to bind endogenous and microbial *trans* ligands mediating inhibitory signals and endocytosis^8, 19, 20, 21^; though *cis* signalling has also been proposed^9, 22, 23^.

Although SIGLEC-5 is known to be expressed in human B-cells^5, 24, 25^, its function in this cell type has not been characterised. Our finding that T-cells expressing CD40L drives SIGLEC-5 expression on B-cells led us to explore a potential role of this B:T-cell regulation. Our data supports the model that stimulated CD4^+^ T-cells display ligands which engage and signal via SIGLEC-5, as determined by SHP1 recruitment. In this regard, it has been widely demonstrated that T-cells undergo a dramatic remodelling on their surface sialoconjugates during development and activation, determining functional outcomes of T-cell interactions with SIGLEC-expressing cells^26, 27, 28, 29^.

Our immunoprecipitation analysis revealed several novel SIGLEC-5 ligand candidates on activated CD4^+^ T-cells, including EBP, a component of the ERC^13^. The ERC is a heterotrimeric complex containing EBP, cathepsin A (CTSA) and neuraminidase 1 (NEU1)^13^. Membrane-bound EBP lacks enzymatic activity, but retains a lectin domain which binds to elastin-derived peptides^12^. Upon binding of EDPs, EBP recruits CTSA and NEU1 and exerts a wide variety of biological functions including chemotaxis, proliferation, and ion influx^13^. These processes are, in part, modulated by the activity of NEU1, which cleaves α2,3, α2,6 and α2,8-sialic acids from cell surface glycoconjugates^30^, therefore suggesting an intrinsic link between the ERC and the bioavailability of SIGLEC ligands^1^. Indeed, our data points to a functional relationship between the ERC and SIGLEC-5. The shedding of EBP leading to the disruption of the ERC, through addition of chondroitin sulphate to T-cells, impaired the ability of SIGLEC-5^+^ B-cells to suppress T-cell IFNγ and IL-17 expression.

Of interest, direct inhibition of EBP with lactose did not alter the capacity of B-cells to suppress IFN-γ□ and IL-17□ T-cells. Lactose targets the galactoside-binding activity of EBP and mainly interferes with canonical recognition of elastin-derived ligands. CHS, by contrast, promotes shedding of surface EBP, and by doing so removes or destabilises the EBP/CTSA-containing complex available for SIGLEC-5 interaction^16, 17^. Thus, SIGLEC-5-dependent suppression does not appear to rely on the lactose-sensitive EBP binding interface, but instead requires the surface availability of EBP or an EBP/CTSA-containing complex on activated T-cells.

The sialidase activity of NEU1 in the plasma membrane is only functional in the presence of the other two ERC components and is essential for ERC-mediated signal transduction^31, 32, 33^. In activated human T-cells, membrane NEU1 has been shown to colocalize with CTLA-4^34^ and to contribute to the desialylation of specific membrane glycoconjugates leading to the release of IFN-γ^35^. Since our immunoprecipitation assays demonstrated that SIGLEC-5 pulled down only EBP and CTSA, but not NEU1, one can speculate that SIGLEC-5 could additionally be playing a role in disassembling the complex, which would render NEU1 inactive and T-cells unresponsive. This direct regulatory mechanism cannot be excluded in the case of SIGLEC-5 and warrants further study.

We describe for the first time that B-cells express SIGLEC-14 only in a secretory form. Following TLR9 stimulation, we observed an early decrease in membrane SIGLEC-5 followed by an increase in secreted SIGLEC-14, while the membrane levels of the latter remained undetectable. This coordinated regulation of SIGLEC-5 and SIGLEC-14 has already been shown in TLR-activated neutrophils and further reflects their paired receptor behaviour^36^. We show that soluble SIGLEC-14 has no detectable impact on SIGLEC-5^+^ B-cell mediated suppression of T-cell IFNγ or IL-17 expression. However, given its higher avidity for sialoconjugates^4^ and based on our immunoprecipitation results showing binding of SIGLEC-14 to T-cell derived ligands, we cannot rule out a role of soluble SIGLEC-14 in other T-cell functions.

In summary, our data describes a new axis in the regulation of B-cell driven T-cell responses. These findings can contribute to expand the arsenal of B-cell tolerance-inducing sialylated polyvalent ligands or liposomal nanoparticles to be used as pharmacologic alternatives for autoimmune diseases^37, 38^. Additionally, the immunosuppressive tumour microenvironment is, in part, driven by tumour cell hypersialylation and overexpression of SIGLEC-5 in tumour-associated macrophages, neutrophils and myeloid-derived suppressor cells^39, 40, 41^. Thus, the SIGLEC-5-ERC axis warrants further investigation as a potential therapeutic target in cancer and autoimmunity.

## Methods

### Recruitment of healthy donors

Peripheral blood samples were collected from healthy donors following informed consent. Ethical approval was obtained from the UCLH Health Service Trust ethics committee, under reference number 14/SC/1200 and West Midlands Coventry and Warwickshire Research Ethics Committee under reference number 17/WM/0332. For experiments, where large numbers of B and T-cells were required, blood cones were purchased from the NHS blood and transplant unit under research ethics number 21/WA/0388. Sample storage complied with requirements of the Data Protection Act 1998.

### SIGLEC-14 deletion polymorphism screening

DNA was extracted from 2×10^6^ peripheral blood mononuclear cells (PBMCs) using the QIAamp® DNA mini kit (Qiagen). The steps from the DNA extraction from blood protocol was followed according to manufacturer’s instructions, but PBMC pellets were used as the starting material and the final DNA was eluted in 100μl of AE buffer. DNA concentration was assessed using a nanodrop spectrophotometer (ThermoFisher).

PCR amplification of SIGLEC-5, SIGLEC-14 and SIGLEC-5/14 fusion specific amplicons was achieved using the previously described primer pairs^5^. PCR amplification was carried out in a 50μl master mix consisting of 25μl 2x iProof^TM^ high fidelity master mix, 2.5μl forward primer (5μM), 2.5μl reverse primer (5μM), 1.5μl DMSO and 250ng DNA template in an 18.5μl DNA/H_2_0 mix. Amplicons were generated using the following PCR parameters: An initial denaturation step of 98°C for 2 mins, followed by 35 cycles of 98°C for 10s, 56°C for 30s and 72°C for 60s. Lastly, a final extension step of 7 mins at 72°C was used. The PCR reactions were assessed for the presence/absence of amplicons at the expected band size on a 1.2% w/v agarose gel in TAE buffer with SYBR^TM^ safe DNA gel stain, using the ChemiDoc^TM^ touch imaging system (Bio-Rad).

### PBMC, serum isolation and cell culture

Peripheral blood (50ml) was collected into heparinised vacutainers for PBMC isolation. PBMCs were isolated from whole blood or cones by Ficoll-based density centrifugation and were cryogenically stored until downstream use.

B-cells, CD4^+^ T-cells, or CD14^+^ classical monocytes were isolated using negative selection kits (STEMCELL Technologies). Cells were cultured with RPMI 1640 containing L-glutamine and NAHCO_3_ (Sigma-Aldrich) supplemented with 10% foetal calf serum (LabTech), 1% penicillin/streptomycin (100U/ml Penicillin+100μg/ml streptomycin; Sigma-Aldrich) at 37°C with 5% CO_2_.

B-cells were incubated at indicated timepoints with either 1μM CpG-C (ODN 2395, Invivogen), 1μM CpG-C with 10μg/ml Affinipure^TM^ goat anti-human IgG and IgM (H+L, Jackson ImmunoResearch), or with 1μg/ml mega CD40L (Enzo Life sciences). In addition, B-cells were cultured with 5μg/ml Brefeldin A (Merck) or 50μM O-propargyl-puromycin (Bio-Techne). CD4^+^ T-cells were cultured with plate-bound 0.5μg/ml anti-CD3 (Hit3a, BD Biosciences) with or without 1μg/ml of soluble anti-CD28 (CD28.2, ThermoFisher), 1μg/ml recombinant human IgG1-Fc (Bio-Techne) or 1μg/ml recombinant human SIGLEC14-Fc (Bio-Techne).

### SIGLEC-5/SIGLEC-14 ELISA

B-cells were cultured at 2.5×10^6^/ml either unstimulated or with 1μM CpG-C, 1μg/ml CD40L, or CpG-C with 10μg/ml anti-IgG+anti-IgM and collected at indicated timepoints. B-cell supernatants were assessed for SIGLEC-5/SIGLEC-14 with a Duoset ELISA kit, according to manufacturer’s instructions (Bio-Techne).

### Flow cytometry and cell sorting

Flow cytometry was performed as previously described^42, 43^. Briefly, ex vivo PBMCs or cultured B and CD4^+^ T-cells were washed with 1xPBS and then stained with LIVE/DEAD fixable blue Dead Cell Stain (Life Technologies) for 20 minutes at room temperature. Cells were then washed with 1xPBS and incubated with FC receptor block (1:10, Miltenyi Biotec) for 10 min at 4°C. For extracellular multi-colour analysis, cells were stained at 4°C for 20min with the following antibodies: CD19 PE-Cy7 (HIB19, BioLegend, 1/50), CD19 BV785 (HIB19, BioLegend, 1/50), CD24 BV421 (ML5, BioLegend, 1/50), CD24 APC (ML5, BioLegend, 1/50), CD25 AF488 (BC96, BioLegend, 1/50), CD27 Percp cy5.5 (M-T271, BioLegend, 1/50), CD38 BV605 (HB-7, BioLegend, 1/50), IgD BUV395 (IA6-2, BD Biosciences, 1/50), IgM BV510 (MHM-88, BioLegend, 1/50), CD3 BV421 (SK7, BioLegend, 1/50), CD4 BV421 (SK7, Biolegend, 1/50), CD4 BV785 (SK3, BioLegend, 1/50), CD4 BUV737 (SK3, BD Biosciences, 1/50), CD14 BV510 (M5E2, BioLegend, 1/50), CD16 BV711 (3G8, BioLegend, 1/50) and SIGLEC-5/14 PE (1A5, BioLegend, 1/50). For panels with more 2 or more Brilliant Violet^TM^ conjugated antibodies, instead of FACS buffer (1x PBS, 1% FCS), the antibody mixes were diluted in BD Horizon^TM^ Brilliant stain buffer (BD Biosciences). Following cell surface staining, cells were then fixed for 20 minutes at 4°C with intracellular fixation buffer (Thermofisher scientific). Cells were washed and left in FACS buffer for later acquisition.

For intracellular cytokine detection, for the last 4h of culture, cells were incubated with complete medium with 50ng/ml PMA (Merck), 250ng/ml ionomycin (Merck) and 5μg/ml brefeldin A (Merck). After the intracellular fixation step, cells were washed twice with 1x permeabilisation buffer (ThermoFisher), before incubation for 40 min at 4°C with the following antibodies: IL-10 APC (JES3-19F1, BioLegend, 1/50), IFNγ BV605 (4S.B3, BioLegend, 1/50), IL-17 BV711 (BL168, BioLegend, 1/50), DAP12 APC (REA900, Miltenyi Biotec, 1/50) and Isotype control APC (REA293, Miltenyi Biotec, 1/50). Lastly cells were washed twice with 1x permeabilisation buffer and then resuspended in FACS buffer for acquisition. Flow cytometric data were collected on an LSR Fortessa (BD Biosciences) using FACS Diva. Data were analysed using Flowjo (TreeStar).

For sorting, cells were isolated using a FACSAria^TM^ Fusion (BD Biosciences) cell sorter, using an 85μm nozzle. Doublets were excluded using combinations of FSC and SSC height and area. Just before acquisition on the sorter, cells were stained with 0.5μg/ml 4,6-diamidino-2-phenylindole (DAPI; Merck). DAPI^+^ dead cells were then excluded. Cells were collected into 1.5ml Eppendorf holder, with an in-built cooling system to keep the cells alive. SIGLEC-5^+^ and SIGLEC-5^-^ B-cells and CD25^-^CD4^+^ T-cells were sorted with a sort purity routinely > 95%.

### In vitro suppression assay

B-cells were isolated by negative selection and stimulated for 72h with 1μM CpG-C. B-cells were stained with CD19 PE-Cy7 (HIB19, BioLegend, 1:25) and SIGLEC-5/14 PE (1A5, BioLegend, 1:25). In addition, on the day of the sort, autologous PBMCs were thawed and stained for CD4 BV785 (SK3, BioLegend, 1:25), CD25 AF488 (BC96, BioLegend, 1:25) and CD14 BV510 (M5E2, BioLegend, 1/25) was included to exclude CD4^+^ monocytes. CD3 staining was avoided to prevent blocking of the antigen for the agonistic anti-CD3 antibody later used.

Sorted SIGLEC-5^+^ and SIGLEC-5^-^ B-cells were co-cultured with CD25^-^CD4^+^ T-cells at a 1:1 ratio and stimulated with 1μM CpG-C and 0.5μg plate-bound anti-CD3 for 72h. Additionally, cells were cultured with/without blocking antibodies against SIGLEC-5/14 (Bio-Techne, 5μg/ml), IL-10/IL-10R (Bio-Techne, 5μg/ml) or the respective IgG1 isotype control (Bio-Techne, 5μg/ml). For the shedding of EBP, cells were incubated with 1mM lactose or 100μM chondroitin sulphate (Sigma-Aldrich). Following stimulation, cells were analysed for CD4^+^ T-cell IFN-γ and IL-17 expression, gated on CD19^-^CD4^+^ T-cells. The percentage suppression of IFN-γ and IL-17 was calculated as a percentage reduction in IFN-γ or IL-17 from CD4^+^ T-cells cultured alone, compared to when B-cell subsets were added to culture.

### Subcellular fractionation and immunoblotting

For immunoblotting, whole cell lysates were collected from 2×10^6^ monocytes, B-cells and CD4^+^ T-cells after lysis with 60μl RIPA buffer (150 mM NaCl, 1.0% IGEPAL^®^ CA-630, 0.5% sodium deoxycholate, 0.1% SDS, 50 mM Tris, pH 8.0, Merck), containing cOmplete^TM^ mini protease inhibitor cocktail (Merck) and phosSTOP^TM^ phosphatase inhibitors (Roche). Cells were incubated with the RIPA mix on ice for 15 mins and then spun at 13,000 for 15 min at 4°C. Supernatants were collected and assessed for protein concentration using a Pierce^TM^ BCA protein assay kit (ThermoFisher). Of the total lysates, 10μg were incubated with NuPAGE^TM^ sample reducing agent and LDS sample buffer (ThermoFisher) and boiled at 70°C for 10 min. Samples were loaded into mPAGE^TM^ 4-12% Bis-Tris precast gels (Merck) alongside a lane with Amersham^TM^ ECL^TM^ full range rainbow marker (Merck), run for 30 minutes in MES SDS running buffer (ThermoFisher) and transferred using trans-blot turbo mini 0.2μm nitrocellulose transfer packs (Bio-Rad). Membranes were blocked with TBS, containing 0.1% Tween-20 and 5% semi-skimmed milk powder (blocking buffer) for 2h at room temperature. Primary antibody incubations were carried out in blocking buffer overnight at 4°C. Secondary antibody incubations were carried out in blocking buffer for 1h at room temperature. Protein bands were detected with the Supersignal^TM^ chemiluminescent substrates and images were acquired on the ChemiDoc^TM^ touch imaging system (Bio-Rad).

For analysis of SIGLEC-5 and SIGLEC-14 in B-cell supernatants by immunoblotting, cells were stimulated for 72h with 1μg/ml CD40L (Invivogen) and the supernatants were concentrated by spinning for 30 mins at 16,000g for 30 minutes in Amicon® 10kDA MWCO ultra centrifuge filter tubes (Merck). For immunoblot analysis, 20μg of concentrated lysate was used for downstream analysis. For immunoblot analysis of non-denatured, non-reduced samples, lysates were not incubated with reducing buffer or boiled, but SDS was present in the MES running buffer.

For subcellular fractionation, 2×10^6^ ex vivo B-cells or 72h anti-CD3 stimulated CD4^+^ T-cells were lysed and separated into fractions using the subcellular protein fractionation kit for cultured cells (ThermoFisher), according to the manufacturer’s instructions. The ratio of buffers used was 100μl:100μl:50μl:50μl:50μl. For downstream immunoblot analysis, half of the lysates were denatured and heated as above, run by SDS-PAGE and transferred to nitrocellulose membranes.

The following primary antibodies were used for immunoblotting: goat anti-human SIGLEC-5/SIGLEC-14 (Bio-Techne, 0.1μg/ml), sheep anti-human beta-galactosidase/elastin binding protein (Bio-Techne, 0.5μg/ml), mouse anti-human cathepsin A (Biolegend, 2μg/ml), mouse anti-human neuraminidase 1 (Bio-Techne, Clone # 688215, 1μg/ml), rabbit anti-human β-actin (Abcam, 0.5μg/ml), goat anti-human CD40 (Bio-Techne, 1μg/ml), goat anti-human Vimentin (Bio-Techne, 1μg/ml), goat anti-human HSP90 (Bio-Techne, 0.5μg/ml), sheep anti-human ST3GAL1 (Bio-Techne, 1μg/ml), goat anti-human SHP1 (Bio-Techne, 0.5μg/ml). The following secondary antibodies were used: donkey anti-goat IgG H+L HRP (ThermoFisher, 0.1μg/ml), donkey anti-sheep IgG H+L HRP (ThermoFisher, 0.1μg/ml), rabbit anti-sheep IgG H+L HRP (ThermoFisher, 0.1μg/ml), goat anti-mouse IgG H+L HRP (ThermoFisher, 0.1 μg/ml), goat anti-rabbit IgG H+L HRP (ThermoFisher, 0.1 μg/ml) and rabbit anti-goat H+L HRP (ThermoFisher, 0.1 μg/ml).

Immunoblots were analysed using ImageLab software v 6.1.0 (Bio-Rad). Ratios of SIGLEC-5 to SIGLEC-14 were determined with the use of a polyclonal antibody that recognises all forms of SIGLEC-5 and SIGLEC-14 on the same blot. Ratios were calculated using the relative quantification feature, selecting the SIGLEC-5 band as the control band for each analysis.

### Direct SIGLEC-5/SIGLEC-14-Fc immunoprecipitation

For the identification and validation of SIGLEC-5 and SIGLEC-14 ligands, 5×10^7^ CD4^+^ T-cells per condition were isolated, cultured for 72h with 0.5μg anti-CD3 (Hit3a, BD Biosciences) and washed twice with 1xPBS. Cells were incubated with 10μg/ml IgG1-Fc (Thr106-Lys330, Biolegend), SIGLEC5-Fc or SIGLEC14-Fc (Bio-Techne) in 1ml of PBS (10μg total recombinant protein) on ice for 1 hour, then washed twice with ice cold PBS. Cell pellets were resuspended in 350μl Pierce^TM^ IP lysis buffer (ThermoFisher), containing cOmplete^TM^ mini protease inhibitor cocktail (Merck) and phosSTOP^TM^ phosphatase inhibitors (Roche) and incubated for 15 min on ice. Lysates were extracted after spinning at 13,000g for 15 mins and protein concentration was quantified using the Pierce^TM^ BCA protein assay kit (ThermoFisher). SIGLEC-5/SIGLEC-14 bound ligands were pulled down using the Dynabeads^TM^ protein G immunoprecipitation kit (ThermoFisher), according to manufacturer’s instructions with some modifications. Briefly, per condition, 500μg of lysate was used for downstream analysis and 20μg of input (SIGLEC-5/SIGLEC-14 tagged) lysates were stored for input controls. 50μl of dynabeads was used per pulldown and washed as per the protocol. The binding antibody was skipped in this experimental setup. Instead, 500μg of lysate was directly incubated with the antibody binding and washing buffer, topped up to 1ml, added to the dynabeads and incubated with light rotation for 30 minutes at room temperature. After the incubation, the post-incubation eluate was stored as a control. The beads were washed, and a non-denaturing elution was carried out, as per the protocol instructions. The input, immunoprecipitates, and post-incubation eluate lysates were denatured and subject to immunoblot analysis as described above in the immunoblot section, or run on a SDS-PAGE for silver staining with the Pierce^TM^ silver stain for mass spectrometry kit (ThermoFisher), according to manufacturer’s instructions. The gel was imaged using the ChemiDoc^TM^ touch imaging system (Bio-Rad) and the image processed in ImageLab (Bio-Rad). Alternatively, for downstream LC-MS/MS analysis, the lysates were left non-denatured. The pulldown lysates were spun dried onto Vivacon® 500 10,000 kDA MWCO filters at 14,000g for 15 min at room temperature and stored at -20°C, until downstream peptide digestion.

### B-T-cell co-culture immunoprecipitations and B-cell immunoprecipitations/pull-down assays

3×10^7^ B-cells were incubated for 72h with 1mg/ml mCD40L or 2.5×10^7^ B-cells and 2.5×10^7^ CD4^+^ T-cells were incubated together for 72h with 0.5μg plate-bound anti-CD3 (Hit3a, BD Biosciences). At the end of the culture, cells were washed twice with ice cold PBS. Cell pellets were resuspended in 350μl Pierce^TM^ IP lysis buffer (ThermoFisher), containing cOmplete^TM^ mini protease inhibitor cocktail (Merck) and phosSTOP^TM^ phosphatase inhibitors (Roche) and incubated for 15 min on ice. Lysates were extracted after spinning at 13,000g for 15 mins and protein concentration was quantified using the Pierce^TM^ BCA protein assay kit (ThermoFisher).

Pulldown of SHP1 recruitment to SIGLEC-5 on B-cells was assessed using the Dynabeads^TM^ protein G immunoprecipitation kit (ThermoFisher). The pulldowns were carried out according to the manufacturer’s instructions, with the following modifications. 50μl of Dynabeads were incubated with 2μg of anti-SIGLEC-5/14 (1A5) for 10 mins at room temperature, with light rotation. 250μg (B-cell IPs) or 500μg (B+T-cell co-culture IPs) were then incubated with the Ab-bead complex for 10 minutes at room temperature, with light rotation. The beads were washed and a denaturing elution was carried out, as per the protocol instructions. The immunoprecipitate lysates were denatured and subject to immunoblot analysis as described above in the immunoblot section. For B-cell IP’s and B:T-cell co-culture IP’s, 30μg of input and post-IP flow through was loaded as a control.

For biotinylation pull-down assays, 1×10^7^ *ex vivo* or 72h mCD40L stimulated B-cells were washed twice with 1xPBS, resuspended at 25M/ml in 1x PBS and then incubation with 5mM EZ-Link Sulfo-NHS-biotin (ThermoFisher) for 30 minutes at room temperature. Cells were washed 3x with 1xPBS+100mM glycine and 1x wash with PBS, before lysates were extracted using IP lysis buffer+ protease inhibitors and Phos-STOP as above. Pulldown of biotinylated proteins were carried out M280 streptavidin Dynabeads^TM^ (ThermoFisher Scientific), according to manufacturer’s instructions. 100μg was loaded for the IP reaction. The resulting lysate and 20% input and post-IP flow through were run on SDS-PAGE gels.

### LC-MS/MS

The pulldown lysates were in filter digested as described previously^44^. Briefly the proteins were spun onto Vivacon® 500 10,000 kDA MWCO filters, were denatured with 8M Urea containing 50mM TCEP-HCl for 1 hour at room temperature and then the filters were spun to dryness at 14,000g. The filters were washed 3x with 200μl of 50mM ammonium bicarbonate, followed by alkylation of free cysteines with 50mM iodoacetamide in 50mM ammonium bicarbonate for 1 hour, at room temperature, in the dark. The denatured proteins were digested overnight at 37°C with 2μg of sequencing grade porcine trypsin (Promega) in 200μl of 50mM ammonium bicarbonate. The peptides were washed through the filter by centrifuging at 14000g and the filter further washed with 0.1% formic acid and 0.1% formic acid/acetonitrile. The eluted peptides were lyophilised prior to LC-MS/MS analysis.

LC-MS/MS analysis was performed on an Orbitrap Eclipse ^TM^ Tribrid mass spectrometer (Thermo scientific). Lyophilised peptides were resuspended in 100μl of 0.1% formic acid and 5μl was injected onto a Pepmax 150cm UHPLC column (Thermo scientific). The peptides were separated over a standard 60 min LC gradient.

Label free quantification (LFQ) analysis was performed using the Peaks studio 11 software suite (Bioinformatics solutions inc). Raw files from triplicate injections of each sample were searched against the human Swissprot database using standard LC-MS/MS search parameters. LFQ was carried out defining the relative protein abundances of proteins with 2 or more unique peptides in the SIGLEC-5-Fc and SIGLEC-14-Fc pulldowns vs the IgG1-Fc control sample.

### scRNA-seq analysis

The scRNA-seq dataset was obtained from Szabo et al^14^, under accession number GSE126030 and analysed using Seurat^45^. The raw count data were normalised using the *NormalizeData()* function, followed by *FindVariableFeatures()* to select the 1000 highest variable genes. The dataset was then scaled using *ScaleData()*. A principal component analysis (PCA) was then performed using *RunPCA(),* and Harmony^46^ was applied to remove the batch effects using the first 10 principal components containing the majority of variance according to the elbow plot. Finally, non-linear dimensionality reduction, nearest-neighbour graph construction, and cluster determination were performed using *RunUMAP()*, *FindNeighbors()*, and *FindClusters()*. T-cell subsets were subsequently extracted using the *subset()* function based on immune lineage markers.

### Statistics

All values are expressed as the mean□±□s.e.m. Data distribution was tested using Shapiro–Wilk normality tests. Statistical significance was determined using unpaired t tests (comparison of two groups), paired t-tests (for matched samples), Mann-Whitney tests (comparison of two groups, non-parametric data), one-way ANOVA (comparison of three or more groups) with Tukey’s multiple comparison post-hoc test or mixed effects analysis with Dunnett’s multiple comparison post-hoc test (multiple comparisons with missing values). Results were considered significant at p ≤ 0.05. Statistical tests were carried out using GraphPad Prism (la Jolla, USA) v10.4.0 (527) Software for Apple Mac.

## Acknowledgements

This work is funded by a Versus Arthritis UK program grant (no. 21140) and from the Innovative Medicines Initiative Joint grant agreement no. 115303, as part of the ABIRISK consortium (Anti-Biopharmaceutical Immunization: prediction and analysis of clinical relevance to minimize the risk) awarded to C. Mauri; and a UCL global engagement grant awarded to C.J.M.Piper. We would like to thank Janani Sivakumaran-Nguyen and Sam Blanchett for cell sorting and flow cytometry expertise and Min Fang (MHRA) for running the LC-MS/MS samples.

## Contributions

C.J.M.P. designed and performed experiments, analysed data and wrote the paper. C.Me performed and analysed the LC MS/MS and critically reviewed the paper. M.L., G.M-G and Z.B. designed and performed experiments. L.N. helped design experiments and critically reviewed the paper. D.C and C.M designed experiments, analysed data and wrote the paper.

## Data availability

The LC-MS/MS data is currently awaiting curation on EMBL-EBI PRIDE accession database.

---

**Extended Data Fig 1.**
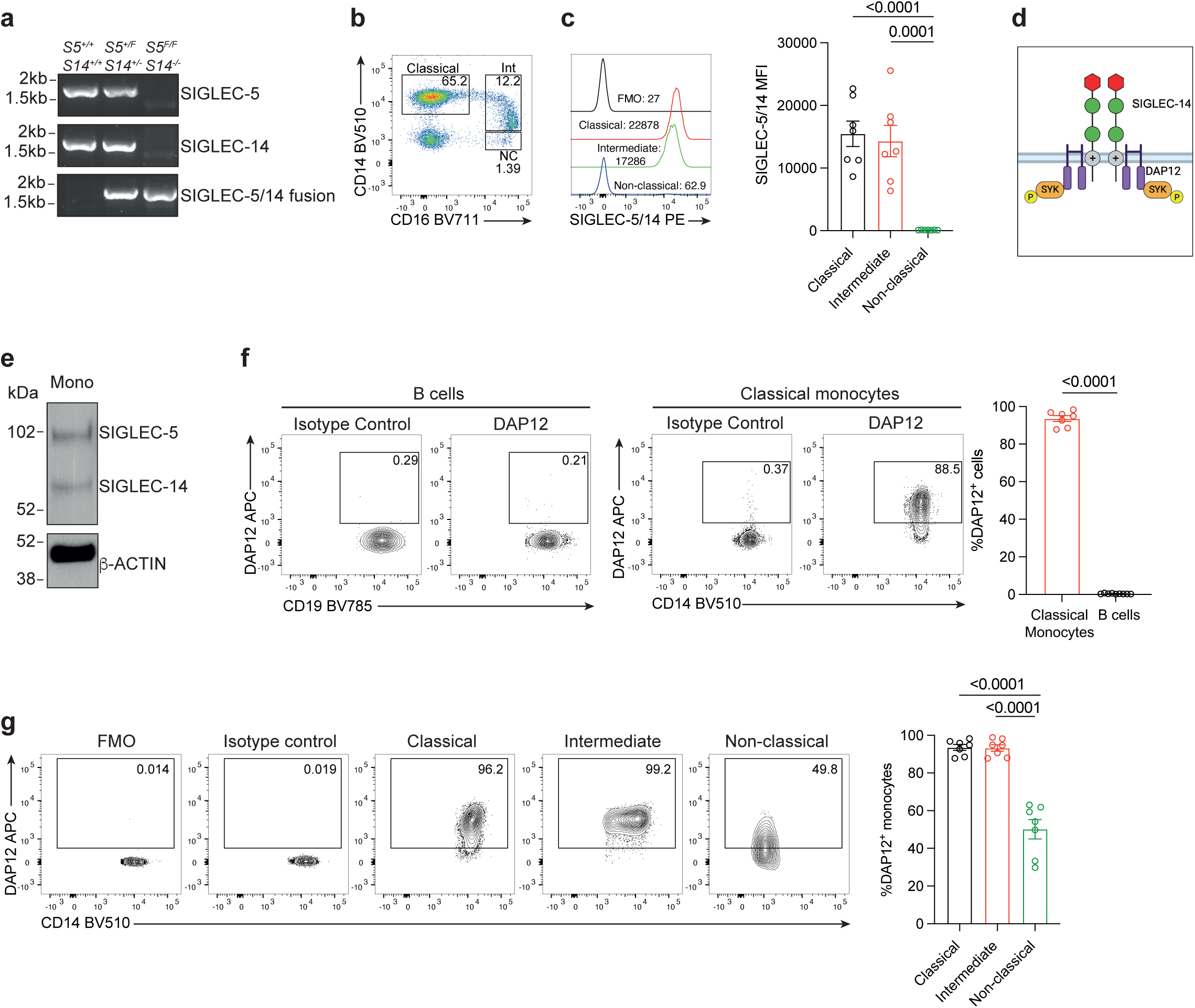
In contrast to monocytes, B-cells do not express DAP12. **a,** Genomic PCR of SIGLEC-5, SIGLEC-14 and SIGLEC-5/14 fusion specific amplicons. **b,** Representative flow cytometry plots showing the gating strategy for the identification of classical, intermediate (int) and non-classical (NC) monocyte subsets. **c,** Representative histograms and bar chart of SIGLEC-5/SIGLEC-14 mean fluorescent intensity (MFI) on monocyte subsets from healthy individuals (n=7). **d,** Schematic illustrating the SIGLEC-14 signalling pathway. **e,** Immunoblots of SIGLEC-5 and SIGLEC-14 expression in monocytes. β-ACTIN was used as a loading control. **f,** Representative flow cytometry plots and bar charts showing the percentage of DAP12^+^ B-cells (left) and DAP12^+^ CD14^+^ classical monocytes (right) in healthy individuals (n=9 and 7 respectively). **g,** Representative flow cytometry plots and bar chart showing the expression of DAP12 in monocyte subsets from healthy individuals (n=7). All data are representative of at least two independent experiments. All values represent the mean ± SEM; Student’s t test for **f**; one-way ANOVA for **c,g**. FMO: fluorescence minus one.

**Extended Data Fig 2.**
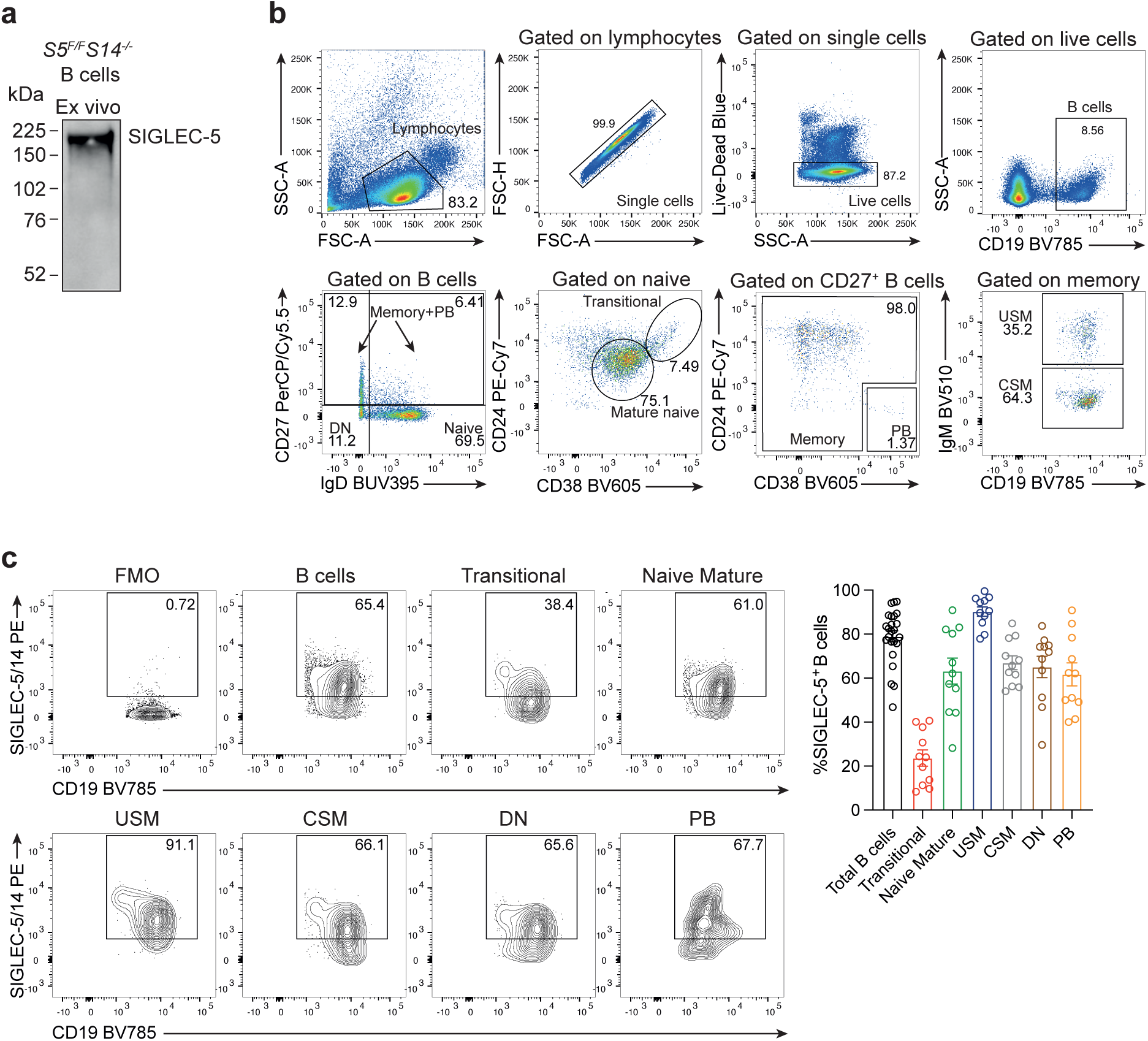
SIGLEC-5 expression increases with B-cell maturity. **a,** Immunoblot of SIGLEC-5 in ex vivo B-cells from SIGLEC-14 null individuals under non-reducing/non-denaturing conditions. **b,** Representative flow cytometry plots showing the gating strategy for the identification of B-cell subsets. DN: Double negative; PB: Plasmablasts; USM: Unswitched memory; CSM: Class-switched memory. **c,** Representative flow cytometry plots and bar chart of the frequency of SIGLEC-5/14^+^ cells in total B-cells (n=23) and B-cell subsets (n=11) from healthy individuals. All data are representative of at least two independent experiments. All values represent the mean ± SEM. FMO: fluorescence minus one.

**Extended Data Fig 3.**
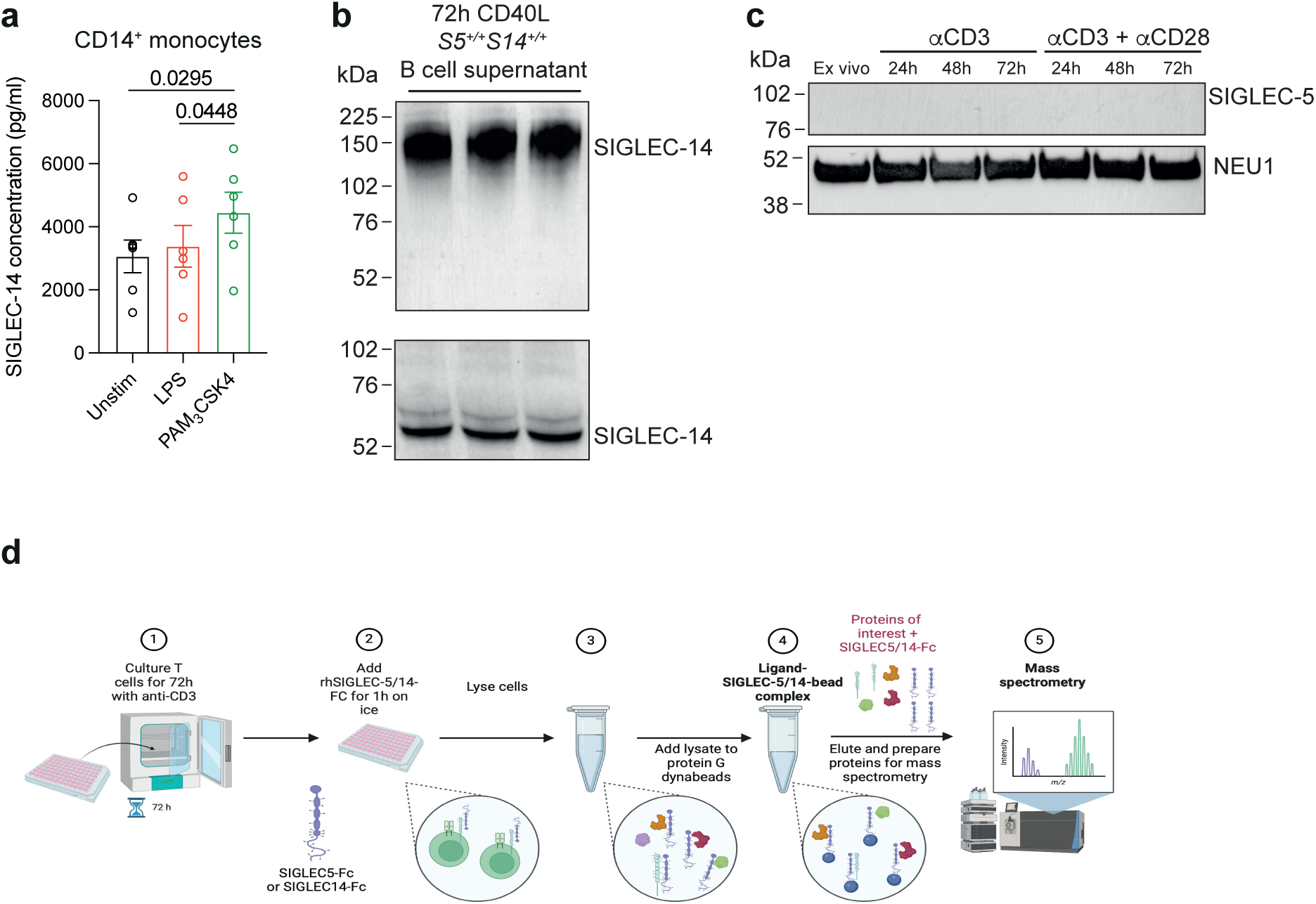
SIGLEC-5 is not expressed in CD4^+^ T-cells. **a,** CD14^+^ monocytes were isolated from healthy PBMC and either left unstimulated or activated with 1μg/ml or 1μg/ml PAM3CSK for 24h. Supernatants were assessed for the levels of SIGLEC-14 by ELISA (n=6). **b,** B-cells were stimulated for 72h with 1μg/ml mCD40L and the supernatants were assessed for SIGLEC-5 and SIGLEC-14 by immunoblot under non-denatured/non-reduced (top blot) and reduced/denatured conditions (bottom blot). **c,** Immunoblots of SIGLEC-5 and NEU1 in 72h αCD3±CD28 stimulated CD4^+^ T-cells. NEU1 was used as a loading control. **d,** Schematic illustrating the experimental design for identifying SIGLEC-5 and SIGLEC-14 ligands on activated CD4^+^ T-cells by mass spectrometry (LC-MS/MS). CD4+ T-cells were tagged with recombinant IgG1-Fc, SIGLEC5-Fc (S5-Fc) or SIGLEC14-Fc (S14-Fc) and associated ligands were immunoprecipitated by protein G bound beads. The resulting lysates were analysed by LC-MS/MS. All data are representative of at least two independent experiments. All values represent the mean ± SEM; one-way ANOVA for **a**. FMO: fluorescence minus one.

**Extended Data Fig 4.**
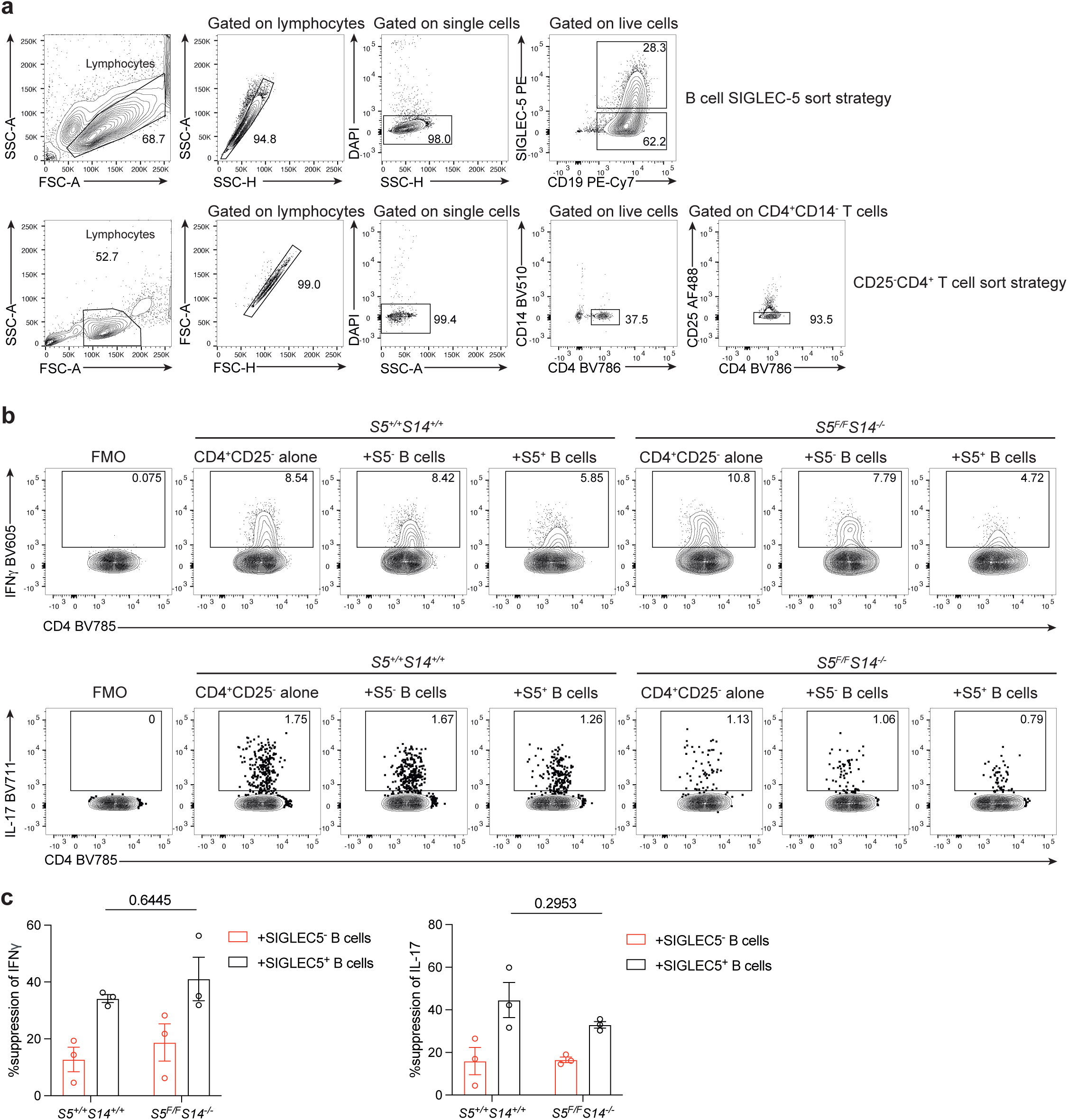
Soluble SIGLEC-14 plays a redundant role in B-cell mediated suppression of T-cell IFNγ and IL-17 expression. **a,** Representative flow cytometry plots showing the gating strategy for sorting SIGLEC-5^+^ and SIGLEC-5^-^ B-cells and CD25^-^CD4^+^ T-cells. **b-c,** B-cells from S5^+/+^S14^+/+^ and S5^F/F^S14^-/-^ individuals were stimulated for 72h with 1μM CpG-C and SIGLEC-5^+^ and SIGLEC-5^-^ B-cells were FACS sorted and co-cultured with CD25^-^CD4^+^T-cells with 0.5μg/ml plate bound anti-CD3 and 1μM CpG-C. **b,** Representative flow cytometry plots and **c,** bar charts showing the percentage suppression of T-cell IFNγ and IL-17 expression after the addition of SIGLEC-5^+^ and SIGLEC-5^-^ subsets (n=3). All data are representative of at least two independent experiments. All values represent the mean ± SEM; Two-way ANOVA for **c** (Šídák multiple comparison post-hoc test). FMO: fluorescence minus one.

**Extended Data Fig 5.**
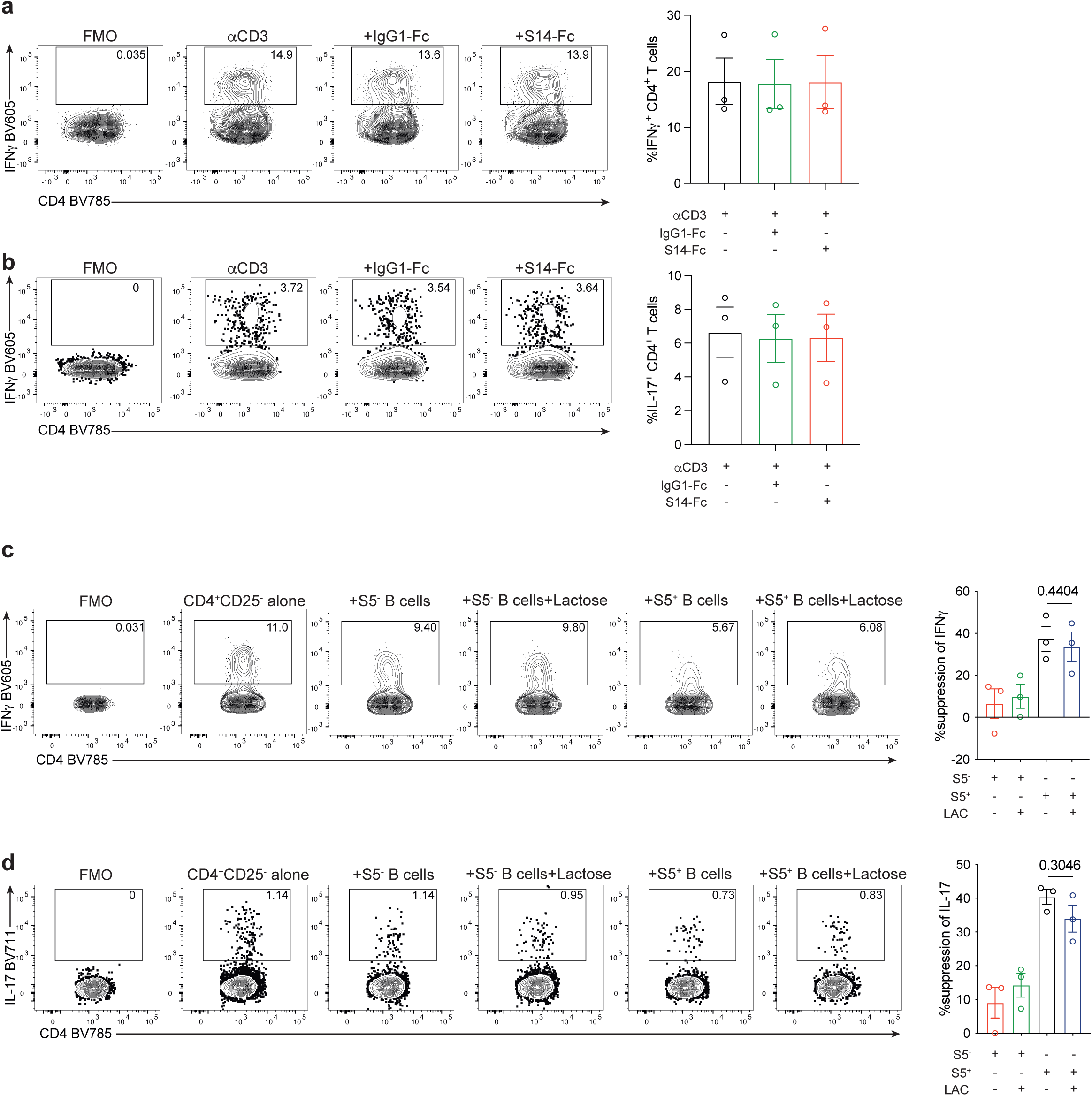
Lactose does not impair SIGLEC-5^+^ B cell suppression of T-cell cytokines. **a-b,** Representative flow cytometry plots and bar charts showing the percentage of **a,** IFN-γ (n=3) and **b,** IL-17 expression (n=3) in CD4^+^ T-cells cultured for 72h with αCD3 either alone or with 1μg/ml of recombinant human IgG1-Fc or S14-Fc. **c-d,** SIGLEC-5^+^ and SIGLEC-5^-^ B-cells were FACS sorted from S5^+/+^S14^+/+^ individuals and co-cultured with anti-CD3+CpG-C stimulated CD4^+^CD25^-^ T-cells with/without 1mM of lactose (LAC). Representative flow cytometry plots and bar charts showing the percentage suppression of T-cell **c,** IFN-γ (n=3) and **d,** IL-17 expression (n=3). Percentage suppression was calculated compared to anti-CD3+CpG-C stimulated CD4^+^CD25^-^ T-cells alone. All data are representative of at least two independent experiments. All values represent the mean ± SEM; one-way ANOVA (Tukey’s multiple comparison post-hoc test) for **c,d**. FMO: fluorescence minus one.

## Notes

### Competing Interest Statement

The authors have declared no competing interest.

