## Supplemental Table 1 for "B-cell SIGLEC-5 engages T-cell components of the elastin receptor complex (ERC) to suppress inflammatory T-cell cytokines"

| Accession | Gene | Significance | Coverage (%) | #Peptides |
|---|---|---|---|---|
| P08670\|VIME_HUMAN | VIM | 12 36 | 61 59 | 37 |
| P02545\|LMNA_HUMAN | LMNA | 4 34 | 6 93 | 3 |
| P10619\|PPGB_HUMAN | CTSA | 25 61 | 8 54 | 4 |
| P16278\|BGAL_HUMAN | GLB1 | 14 9 | 11 67 | 6 |
| P20963\|CD3Z_HUMAN | CD247 | 1 62 | 50 61 | 7 |
| P07339\|CATD_HUMAN | CTSD | 0 38 | 6 55 | 3 |
| P63173\|RL38_HUMAN | RPL38 | 2 83 | 32 86 | 2 |
| Q8WZ42\|TITIN_HUMAN | TTN | 1 97 | 0 13 | 3 |
| P07355\|ANXA2_HUMAN | ANXA2 | 15 14 | 23 3 | 6 |
| P20930\|FILA_HUMAN | FLG | 0 93 | 1 97 | 3 |
| P07766\|CD3E_HUMAN | CD3E | 0 96 | 49 76 | 7 |
| P01848\|TRAC_HUMAN | TRAC | 2 65 | 15 | 2 |
| P0DTU3\|TRAR2_HUMAN | TRA | 2 65 | 7 64 | 2 |
| P04234\|CD3D_HUMAN | CD3D | 0 5 | 18 71 | 3 |
| P09693\|CD3G_HUMAN | CD3G | 0 81 | 12 64 | 2 |
| P04406\|G3P_HUMAN | GAPDH | 12 51 | 21 19 | 7 |
| Q9BY77\|PDIP3_HUMAN | POLDI | 1 57 | 4 75 | 2 |
| P27816\|MAP4_HUMAN | MAP4 | 0 93 | 5 38 | 6 |
| Q08188\|TGM3_HUMAN | TGM3 | 6 14 | 3 75 | 3 |
| P62987\|RL40_HUMAN | UBA52 | 5 24 | 13 28 | 3 |
| P38646\|GRP75_HUMAN | HSPA9 | 0 31 | 3 98 | 4 |
| P05089\|ARGI1_HUMAN | ARG1 | 4 97 | 9 01 | 3 |
| B0I1T2\|MYO1G_HUMAN | MYO1G | 6 8 | 9 72 | 7 |
| P35579\|MYH9_HUMAN | MYH9 | 1 16 | 3 27 | 6 |
| P01876\|IGHA1_HUMAN | IGHA1 | 10 69 | 12 81 | 6 |
| P81605\|DCD_HUMAN | DCD | 3 89 | 56 36 | 12 |
| P31151\|S10A7_HUMAN | S100A | 2 68 | 30 69 | 2 |
| Q02383\|SEMG2_HUMAN | SEMG2 | 1 3 | 5 33 | 3 |
| A6NMY6\|AXA2L_HUMAN | ANXA2 | 16 38 | 13 57 | 4 |
| Q15517\|CDSN_HUMAN | CDSN | 4 72 | 8 32 | 3 |
| Q5D862\|FILA2_HUMAN | FLG2 | 4 57 | 2 68 | 5 |
| P01040\|CYTA_HUMAN | CSTA | 4 98 | 68 37 | 6 |
| P68104\|EF1A1_HUMAN | EEF1A | 6 24 | 4 76 | 5 |
| Q5VTE0\|EF1A3_HUMAN | EEF1A | 6 24 | 4 76 | 5 |
| Q9NZT1\|CALL5_HUMAN | CALML | 5 84 | 50 | 8 |
| P10599\|THIO_HUMAN | TXN | 7 41 | 20 95 | 2 |
| P02768\|ALBU_HUMAN | ALB | 2 43 | 26 44 | 19 |
| Q5T749\|KPRP_HUMAN | KPRP | 1 93 | 37 65 | 14 |
| P14923\|PLAK_HUMAN | JUP | 7 37 | 34 5 | 18 |
| P15924\|DESP_HUMAN | DSP | 3 35 | 17 24 | 44 |
| P06702\|S10A9_HUMAN | S100A | 8 74 | 43 86 | 4 |
| Q02413\|DSG1_HUMAN | DSG1 | 10 31 | 20 88 | 14 |
| Q01469\|FABP5_HUMAN | FABP5 | 7 22 | 35 56 | 4 |
| Q86YZ3\|HORN_HUMAN | HRNR | 5 63 | 10 39 | 20 |
| P05109\|S10A8_HUMAN | S100A | 10 36 | 31 18 | 3 |
| P62805\|H4_HUMAN | H4C16 | 4 52 | 31 07 | 3 |

| | | | | | | |
|---|---|---|---|---|---|---|
| Q08554\|DSC1_HUMAN | DSC1 | | 10 | 8 | 17 | 6 |
| P47929\|LEG7_HUMAN | LGALS | 9 | 48 | 27 | 94 | 3 |
| Q15149\|PLEC_HUMAN | PLEC | 10 | 63 | 7 | 84 | 34 |

| #Unique | Group Profile (Ratio) | | | Avg. Mass |
|---|---|---|---|---|
| | IgG | Siglec-5 | Siglec-14 | |
| 28 | 1 | 14 19 | 46 83 | 53652 |
| 3 | 1 | 6 17 | 1 03 | 74139 |
| 4 | 1 | 2 3 | 6 17 | 54466 |
| 6 | 1 | 1 45 | 4 72 | 76075 |
| 7 | 1 | 1 44 | 1 16 | 18696 |
| 3 | 1 | 1 38 | 1 07 | 44552 |
| 2 | 1 | 1 33 | 0 97 | 8218 |
| 3 | 1 | 1 31 | 0 83 | 3816030 |
| 6 | 1 | 1 3 | 1 82 | 38604 |
| 3 | 1 | 1 28 | 0 76 | 435170 |
| 7 | 1 | 1 27 | 1 07 | 23147 |
| 2 | 1 | 1 26 | 0 91 | 15717 |
| 2 | 1 | 1 26 | 0 91 | 30989 |
| 3 | 1 | 1 23 | 1 07 | 18930 |
| 2 | 1 | 1 22 | 1 06 | 20469 |
| 7 | 1 | 1 17 | 0 35 | 36053 |
| 2 | 1 | 1 16 | 1 01 | 46089 |
| 6 | 1 | 0 99 | 0 95 | 121005 |
| 3 | 1 | 0 98 | 0 78 | 76632 |
| 3 | 1 | 0 95 | 0 71 | 14728 |
| 3 | 1 | 0 9 | 0 61 | 73680 |
| 3 | 1 | 0 79 | 1 94 | 34735 |
| 7 | 1 | 0 78 | 0 29 | 116442 |
| 5 | 1 | 0 73 | 0 61 | 226532 |
| 3 | 1 | 0 7 | 2 9 | 42849 |
| 12 | 1 | 0 7 | 1 27 | 11284 |
| 2 | 1 | 0 67 | 1 14 | 11471 |
| 3 | 1 | 0 66 | 3 04 | 65444 |
| 4 | 1 | 0 65 | 1 59 | 38659 |
| 3 | 1 | 0 6 | 1 06 | 51607 |
| 5 | 1 | 0 57 | 1 2 | 248073 |
| 6 | 1 | 0 55 | 1 02 | 11006 |
| 3 | 1 | 0 53 | 0 76 | 50141 |
| 3 | 1 | 0 53 | 0 76 | 50185 |
| 8 | 1 | 0 51 | 1 36 | 15893 |
| 2 | 1 | 0 5 | 1 07 | 11737 |
| 16 | 1 | 0 49 | 0 97 | 69367 |
| 14 | 1 | 0 47 | 1 74 | 64136 |
| 17 | 1 | 0 45 | 1 48 | 81745 |
| 42 | 1 | 0 44 | 1 49 | 331774 |
| 4 | 1 | 0 32 | 1 17 | 13242 |
| 14 | 1 | 0 31 | 0 97 | 113748 |
| 4 | 1 | 0 29 | 1 06 | 15164 |
| 20 | 1 | 0 29 | 0 85 | 282390 |
| 3 | 1 | 0 27 | 0 8 | 10835 |
| 3 | 1 | 0 24 | 0 28 | 11367 |

| | | | | | | |
|---|---|---|---|---|---|---|
| 6 | 1 | 0 | 22 | 1 | 24 | 99987 |
| 3 | 1 | 0 | 2 | 0 | 2 | 15075 |
| 30 | 1 | 0 | 12 | 1 | 34 | 531791 |

Description

Vimentin OS=Homo sapiens OX=9606 GN=VIM PE=1 SV=4
Prelamin-A/C OS=Homo sapiens OX=9606 GN=LMNA PE=1 SV=1
Lysosomal protective protein OS=Homo sapiens OX=9606 GN=CTSA PE=1 SV=2
Beta-galactosidase OS=Homo sapiens OX=9606 GN=GLB1 PE=1 SV=2
T-cell surface glycoprotein CD3 zeta chain OS=Homo sapiens OX=9606 GN=CD247 PE=1 SV=2
Cathepsin D OS=Homo sapiens OX=9606 GN=CTSD PE=1 SV=1
Large ribosomal subunit protein eL38 OS=Homo sapiens OX=9606 GN=RPL38 PE=1 SV=2
Titin OS=Homo sapiens OX=9606 GN=TTN PE=1 SV=4
Annexin A2 OS=Homo sapiens OX=9606 GN=ANXA2 PE=1 SV=2
Filaggrin OS=Homo sapiens OX=9606 GN=FLG PE=1 SV=3
T-cell surface glycoprotein CD3 epsilon chain OS=Homo sapiens OX=9606 GN=CD3E PE=1 SV=2
T cell receptor alpha chain constant OS=Homo sapiens OX=9606 GN=TRAC PE=1 SV=2
T cell receptor alpha chain MC.7.G5 OS=Homo sapiens OX=9606 GN=TRA PE=1 SV=1
T-cell surface glycoprotein CD3 delta chain OS=Homo sapiens OX=9606 GN=CD3D PE=1 SV=1
T-cell surface glycoprotein CD3 gamma chain OS=Homo sapiens OX=9606 GN=CD3G PE=1 SV=1
Glyceraldehyde-3-phosphate dehydrogenase OS=Homo sapiens OX=9606 GN=GAPDH PE=1 SV=3
Polymerase delta-interacting protein 3 OS=Homo sapiens OX=9606 GN=POLDIP3 PE=1 SV=2
Microtubule-associated protein 4 OS=Homo sapiens OX=9606 GN=MAP4 PE=1 SV=3
Protein-glutamine gamma-glutamyltransferase E OS=Homo sapiens OX=9606 GN=TGM3 PE=1 SV=4
Ubiquitin-ribosomal protein eL40 fusion protein OS=Homo sapiens OX=9606 GN=UBA52 PE=1 SV=2
Stress-70 protein, mitochondrial OS=Homo sapiens OX=9606 GN=HSPA9 PE=1 SV=2
Arginase-1 OS=Homo sapiens OX=9606 GN=ARG1 PE=1 SV=2
Unconventional myosin-Ig OS=Homo sapiens OX=9606 GN=MYO1G PE=1 SV=2
Myosin-9 OS=Homo sapiens OX=9606 GN=MYH9 PE=1 SV=4
Immunoglobulin heavy constant alpha 1 OS=Homo sapiens OX=9606 GN=IGHA1 PE=1 SV=3
Dermcidin OS=Homo sapiens OX=9606 GN=DCD PE=1 SV=2
Protein S100-A7 OS=Homo sapiens OX=9606 GN=S100A7 PE=1 SV=4
Semenogelin-2 OS=Homo sapiens OX=9606 GN=SEMG2 PE=1 SV=1
Putative annexin A2-like protein OS=Homo sapiens OX=9606 GN=ANXA2P2 PE=5 SV=2
Corneodesmosin OS=Homo sapiens OX=9606 GN=CDSN PE=1 SV=4
Filaggrin-2 OS=Homo sapiens OX=9606 GN=FLG2 PE=1 SV=1
Cystatin-A OS=Homo sapiens OX=9606 GN=CSTA PE=1 SV=1
Elongation factor 1-alpha 1 OS=Homo sapiens OX=9606 GN=EEF1A1 PE=1 SV=1
Putative elongation factor 1-alpha-like 3 OS=Homo sapiens OX=9606 GN=EEF1A1P5 PE=5 SV=1
Calmodulin-like protein 5 OS=Homo sapiens OX=9606 GN=CALML5 PE=1 SV=2
Thioredoxin OS=Homo sapiens OX=9606 GN=TXN PE=1 SV=3
Albumin OS=Homo sapiens OX=9606 GN=ALB PE=1 SV=2
Keratinocyte proline-rich protein OS=Homo sapiens OX=9606 GN=KPRP PE=1 SV=1
Junction plakoglobin OS=Homo sapiens OX=9606 GN=JUP PE=1 SV=3
Desmoplakin OS=Homo sapiens OX=9606 GN=DSP PE=1 SV=3
Protein S100-A9 OS=Homo sapiens OX=9606 GN=S100A9 PE=1 SV=1
Desmoglein-1 OS=Homo sapiens OX=9606 GN=DSG1 PE=1 SV=2
Fatty acid-binding protein 5 OS=Homo sapiens OX=9606 GN=FABP5 PE=1 SV=3
Hornerin OS=Homo sapiens OX=9606 GN=HRNR PE=1 SV=2
Protein S100-A8 OS=Homo sapiens OX=9606 GN=S100A8 PE=1 SV=1
Histone H4 OS=Homo sapiens OX=9606 GN=H4C16 PE=1 SV=2

Desmocollin-1 OS=Homo sapiens OX=9606 GN=DSC1 PE=1 SV=2
Galectin-7 OS=Homo sapiens OX=9606 GN=LGALS7B PE=1 SV=2
Plectin OS=Homo sapiens OX=9606 GN=PLEC PE=1 SV=3
